# Thermodynamically consistent determination of free energies and rates in kinetic cycle models

**DOI:** 10.1101/2023.04.08.536126

**Authors:** Ian M. Kenney, Oliver Beckstein

## Abstract

Kinetic and thermodynamic models of biological systems are commonly used to connect microscopic features to system function in a bottom-up multiscale approach. The parameters of such models—free energy differences for equilibrium properties and in general rates for equilibrium and out-of-equilibrium observables—have to be measured by different experiments or calculated from multiple computer simulations. All such parameters necessarily come with uncertainties so that when they are naively combined in a full model of the process of interest, they will generally violate fundamental statistical mechanical equalities, namely detailed balance and an equality of forward/backward rate products in cycles due to T. Hill. If left uncorrected, such models can produce arbitrary outputs that are physically inconsistent. Here we develop a maximum likelihood approach (named *multibind*) based on the so-called potential graph to combine kinetic or thermodynamic measurements to yield state resolved models that are thermodynamically consistent while being most consistent with the provided data and their uncertainties. We demonstrate the approach with two theoretical models, a generic two-proton binding site and a simplified model of a sodium/proton antiporter. We also describe an algorithm to use the *multibind* approach to solve the inverse problem of determining microscopic quantities from macroscopic measurements and as an example we predict the microscopic p*K*_a_s and protonation states of a small organic molecule from 1D NMR data. The *multibind* approach is applicable to any thermodynamic or kinetic model that describes a system as transitions between well-defined states with associated free energy differences or rates between these states. A Python package `multibind`, which implements the approach described here, is made publicly available under the MIT Open Source license.

**WHY IT MATTERS:** The increase in computational efficiency and rapid advances in methodology for quantitative free energy and rate calculations has allowed for the construction of increasingly complex thermodynamic or kinetic “bottom-up” models of chemical and biological processes. These multi-scale models serve as a framework for analyzing aspects of cellular function in terms of microscopic, molecular properties and provide an opportunity to connect molecular mechanisms to cellular function. The underlying model parameters—free energy differences or rates—are constrained by thermodynamic identities over cycles of states but these identities are not necessarily obeyed during model construction, thus potentially leading to inconsistent models. We address these inconsistencies through the use of a maximum likelihood approach for free energies and rates to adjust the model parameters in such a way that they are maximally consistent with the input parameters and exactly fulfill the thermodynamic cycle constraints. This approach enables formulation of thermodynamically consistent multi-scale models from simulated or experimental measurements.

## INTRODUCTION

Living cells perform a multitude of functions such as cell signaling, reaction catalysis, gene expression, DNA replication, or transmembrane transport (1). At the molecular level, these cellular processes can be represented as kinetic networks (2–13) that consist of distinct states of molecules and the reactions that interconvert between the states. In a kinetic network (or kinetic graph), each reaction is quantified by its forward and reverse rate, and so a set of master equations relates the change in the population of each state to fluxes between states via mass-action kinetics (14). When parameterized correctly, these networks provide a powerful framework for analyzing non-equilibrium systems in a variety of external conditions; for example, macroscopic observables can be calculated from microscopic quantities in a bottom-up multiscale approach (6, 7, 9, 13) and pathways within the network can be identified to gain insights into underlying molecular mechanisms (4–7, 15).

However, the construction of these models requires some care to satisfy a number of fundamental physical constraints, as we will outline briefly: Consider a kinetic graph consisting of one or more full cycles, such as the one shown in Figure 1. Each reaction is modeled with mass-action kinetics and restricted to simple elementary reactions (16), i.e., conformational changes of a molecule A between states *i* and *j*, A_*i*_ ⇌ A _*j*_, or binding of a single ligand, *A* + *X* ⇌ *A* : *X*, so that the model can be interpreted in terms of molecular events. The values of external parameters responsible for thermodynamic driving forces such as concentrations [X] or electrostatic potentials Φ are also part of the model so it is convenient to represent all reactions as first order reactions by, if necessary, augmenting the intrinsic rate constants 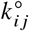 so that they become dependent on the external parameters (e.g., *k*_*ij*_([X]) or *k*_*ij*_ (Φ)) (2). Depending on the external parameters, the kinetic model describes a system in equilibrium or out of equilibrium. Regardless of the overall state of the system, the product of the forward 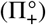 and backward 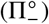 intrinsic rates of *all* closed paths must satisfy Hill’s kinetic cycle closure condition (2)

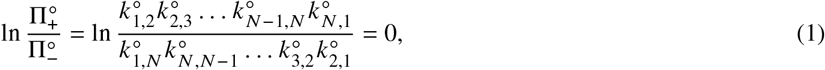

which is ultimately due to the fact that if the system were to be in equilibrium, only the intrinsic rates would matter and then the relationship follows from detailed balance; however, the intrinsic rates do not depend on the external parameters and therefore the same relationship must also hold in all non-equilibrium states.

**Figure 1:**
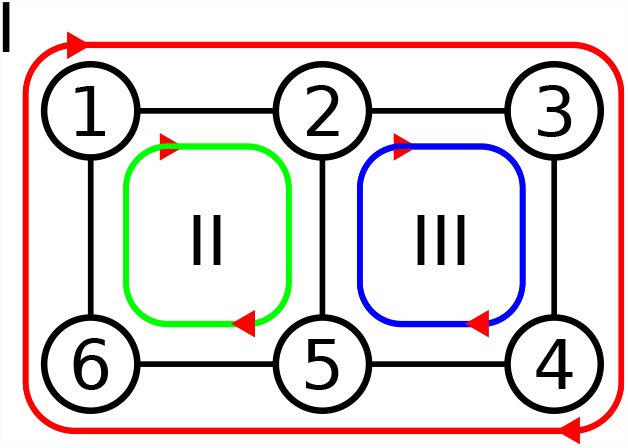
Closed cycles inside a kinetic graph. Each cycle is characterized by the forward and backward rates between each of its states. For instance, the green cycle (2), has forward and backward rates *k*_1,2_, *k*_2,5_, *k*_5,6_, *k*_6,1_, and *k*_1,6_, *k*_6,5_, *k*_5,2_, *k*_2,1_, respectively. All closed cycles must satisfy the kinetic cycle closure condition, Equation 1.

Equation 1 shows that the rates are not independent, with every possible cycle adding additional interdependencies. Therefore, setting up a kinetic network with rates obtained from experiments or simulations may easily lead to violations of the cycle closure condition with the consequence that non-equilibrium systems will have arbitrary, unphysical driving forces embedded into them while equilibrium systems will violate detailed balance and the path-independence of the free energies; thus any predictions made from such a thermodynamically inconsistent model will be incorrect.

In the special case of equilibrium, the kinetic network has to obey detailed balance and instead of forward and backward rates, free energy differences Δ*G*_*ij*_ = *G*_*j*_ − *G*_*i*_ between states completely describe the system. The description can be simplified further by focusing on the thermodynamic free energies of the states *G*_*i*_ themselves. These state free energies are thermodynamic potentials, i.e., differences in free energies are path independent. In particular, the sum of free energy differences between states along any cycle *C* has to vanish; Equation 2 is generally referred to as the equilibrium cycle closure condition (or Wegscheider condition (17)). Because potential functions are used in the study of Markov random fields (also known as undirected graphical models) (18) we call the equilibrium kinetic graph with the state free energies *G*_*i*_ assigned to each node *i* the *potential graph*. Potential graphs satisfy cycle closure by construction since the sum of free energy differences in a closed path always vanishes (because in the sum, each state free energy appears twice but with opposite signs). However, the state free energies themselves are rarely directly available. Instead, free energy differences 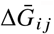 (such as binding free energies) are measured or calculated from simulations. When these free energy differences are obtained from different sources or experiments or simply have statistical errors then these free energy differences cannot be combined naively in a full model because the free energy of each state becomes dependent on the choice of reference state as well as the path taken from that reference state (19–24). This problem has been recognized, in particular in the area of binding free energy calculations, and the standard approach is to solve for the potential graph with a maximum likelihood estimator with the 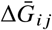 as inputs (21, 22, 24). The resulting potential graph contains all information to compute equilibrium properties based on the graph’s partition function.

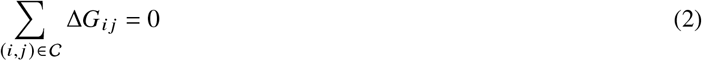

The potential graph approach also suggests how to obtain thermodynamically consistent rates. In the systems biology community an energy-based formalism has been used where rates are modeled as an Arrhenius-like activated process 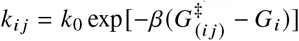. Models are formulated in terms of state free energies *G*_*i*_, transition state free energies 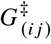 and frequency prefactors *k*_0_ (25–27); this approach has also been used to study kinetic networks of the type introduced above for transporter proteins (7, 15). A somewhat related problem is inferring rate matrix (or transition matrix) parameters from sets of simulations. It has been recognized for a long time that detailed balance can be built into a rate matrix of a kinetic model to infer the model parameters via Bayesian analysis from molecular dynamics simulations (28). Similarly, estimating a reversible transition matrix is crucial for constructing thermodynamically correct Markov state models from simulations (29, 30). Here we develop an alternative approach that directly uses the rate constants and their measurement uncertainties. It has the advantage that rate constants are adjusted towards consistency in forward/backward pairs so that relatively larger uncertainties allow relatively larger changes. Our primary goal was to provide a general computational method to set up thermodynamically consistent equilibrium and non-equilibrium kinetic graphs for molecular systems, given estimates for either free energy differences or rate constants. We named our approach *multibind* and provide and open source software implementation in Python.

The rest of the paper is organized as follows: We first outline the underlying theory for both equilibrium and non-equilibrium kinetic models built upon potential graphs. We then show how the potential graph’s state free energies (together with error estimates) can be calculated from the input free energy differences. Based on the potential graph (and following Hill’s reasoning for Equation 1) we derive a projection approach to find the rates that are thermodynamically consistent. We also provide a simple Monte Carlo algorithm to solve the inverse problem of inferring microscopic rates of a cyclic kinetic graph from macroscopic measurements. We apply our method to three example systems. As a first pedagogical example we demonstrate how the potential graph formalism is able to capture cooperative and anti-cooperative behavior in a simple model of a two proton binding site system. The simple model is then extended to a three proton binding model for the organic molecule diethylenetriaminepentaacetic acid (DTPA) where the microscopic p*K*_a_s are determined from experimental 1D NMR data with the inverse approach. The resulting microscopic model clearly displays cooperative behavior that explains why site-specific protonation curves show strongly non-monotonic behavior as function of pH. As the final example we analyze a simple model of a sodium/proton antiporter secondary active transporter in equilibrium and out of equilibrium with an electrochemical driving force.

## METHODS

We begin by describing how we define states, which form the basis of kinetic models, and the relationship between state free energies, in particular with a view towards the calculation of macroscopic observables from microscopic models. We then describe an approach to solve for the state free energies from equilibrium free energy differences and extend it to also solve for thermodynamically consistent rate constants.

### State space separation and free energies

A classical *N*-particle system with Hamiltonian ℋ in the canonical ensemble is characterized in equilibrium by its partition function *Q* ∝ ∬ *dp dq e*^−*β*ℋ(*p,q*)^ (or equivalently its Helmholtz free energy *βF* = − ln *Q* with *β* = 1/*kT*) where the 3*N* generalized momenta *p* and positions *q* are considered a point in phase space and characterize the microscopic state (31). In order to make clear the relationship between arbitrary collections of microstates, we separate phase space into *M* discrete states by indicator functions *I*_*i*_ (*p, q*) that equal one when a phase space point is within some state definition (such as “ligand bound” or “protein in conformation A”) and zero when outside of this boundary (e.g., “ligand not bound” or “protein in conformation B”). Because the indicator functions partition all of phase space, their sum must be the identity,

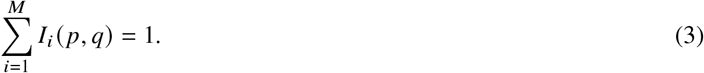

Inserting the sum of indicator functions into the partition function does not change the partition function but decomposes it into a sum of partition functions for the subsystems, *Q*_*i*_,

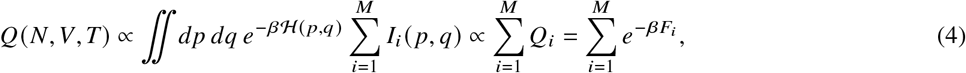

each with a corresponding state Helmholtz free energy, *F*_*i*_. In the isothermal-isobaric ensemble the equivalent relationship holds for the state Gibbs free energies,

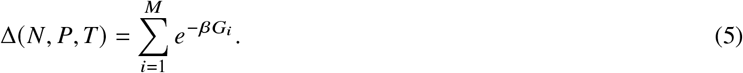

The separation in state space can be made arbitrary and thus provides a general prescription for how to lump states together. In the following we consider a (molecular) *microstate* to be a specific chemical configuration of the system such as a tautomer of a molecule or a single conformer. Such a microstate is described by a finite volume in phase space and corresponding free energy *G*_*i*_. The probability to observe the system in this microstate is

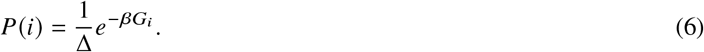

As an example consider a diprotic acid, a molecule with two titratable groups that each can bind a proton. It has four microstates: one each with either zero or two protons bound, and two with one proton bound.

Multiple molecular microstates may be lumped together to describe a *macrostate*—for instance, to describe an experimentally observable state of the system. Each macrostate contains one or more microstates and each microstate is only part of a single macrostate. For the example of the diprotic acid, we might consider the total number of protons bound as the macrostate, resulting in three macrostates. Because a macrostate *s*_*α*_ is a union of microstates, the same considerations as for microstates apply, and the probability to observe the macrostate is

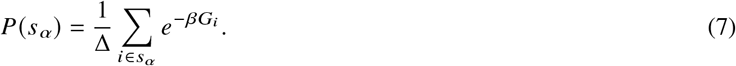

The *macrostate free energy difference*

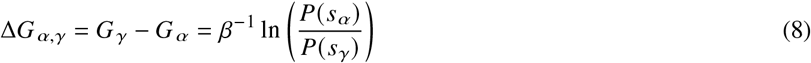

quantifies the relative probability of one macrostate *α* compared to another macrostate *γ*. In general, free energy differences relative to a common reference value are sufficient to completely determine the equilibrium behavior of a system. In practice, experiments and simulations often only provide free energy differences between states and we are left with the problem to find the corresponding state free energies.

It should be noted that for sufficiently complicated systems, it is likely that particular regions of phase space will not be included. The consequences of failing to include states depends on the contribution of that state to the overall partition function through its free energy. For instance, not including short-lived states, such as short timescale transition states connecting two metastable states, will show little impact on model predictions. On the other hand, if a major state is omitted from the model, predictions of state probabilities will certainly be incorrect, which will invalidate many downstream analyses.

### Maximum likelihood estimate of state free energies from free energy differences

The free energies provided by many methods, such as free energy perturbation methods, are given in terms of the free energy difference between two states, rather than the free energies of each. Since the free energy is a state function, the differences can be summed along any path from a common reference state to yield all other state free energies (Equation 2). However, summing along a path only yields consistent state free energies if the the free energy differences are exact. In reality, the differences between pairs of states are usually collected independently of one another and therefore come with their own errors. Thus, state free energies generated by summing along paths become path-dependent even though the free energy function has to be path-independent. Instead, a global approach is needed that considers all state free energies and all free energy differences together. As outlined in our earlier work (24), we calculate state free energies, { *G*_*i*_}, with a global maximum-likelihood approach that uses the measured free energy differences and their associated errors between states *i* and *j*, 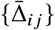 and 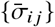, as inputs. Essentially the same idea has been used multiple times in the field of binding free energy calculations (21, 32, 33). Because the maximum likelihood approach is the basis for all our following work, we describe it here in more detail than in Selwa et al. (24) and extend it to also include error estimates. Under the assumption that free energy measurements are Gaussian distributed, the likelihood function is

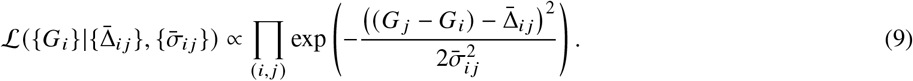

where the product is over all edges (*i, j*) between states *i* and *j* for which there are data. Maximizing the likelihood function with respect to the {*G*_*i*_} yields a set of thermodynamically consistent state free energies *G*_*i*_, up to an additive constant. Because these {*G*_*i*_} form a potential graph, any path dependence is eliminated by construction while simultaneously estimating the set of thermodynamically consistent free energies that are optimally compatible with the input data. In the following we assume that an arbitrary state is chosen as a reference and assigned a free energy of zero in lieu of a known absolute free energy. All other state free energies are then calculated relative to the arbitrary reference state, and for simplicity we also denote these state free energies as {*G*_*i*_}. This approach does not affect any computable observables because a common offset in the state free energies cancels in all expressions involving the partition function; in other words, the physics of the problem is determined by the differences in free energies between states. As shown in Appendix A.1, the error estimate

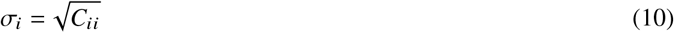

for the state free energy *G*_*i*_ is the square root of the diagonal element of the covariance matrix for the likelihood estimator (32, 33). The covariance matrix is the inverse C = I^−1^ of the Fisher information matrix

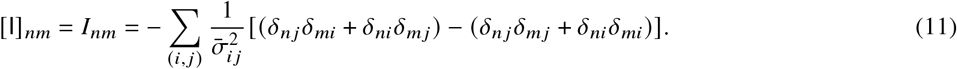

The *δ*_*ij*_ are Kronecker deltas that select the relevant inverse variances of the input free energies for connections between states *i* and *j*. In practice, C is robustly calculated as the Moore-Penrose pseudoinverse of I, C = I^+^, using singular value decomposition (34).

### Thermodynamically consistent rate constants

Kinetic networks introduce time and dynamics to an otherwise static view of a system. They also provide the framework to easily move outside of equilibrium, as they are not constrained by a partition function. In a kinetic network, the focus is on the connecting processes rather than the states themselves. The thermodynamic driving potential χ around any cycle within this network is related to the ratio of products of the forward (Π_+_) and reverse (Π_−_) rates around the cycle (2),

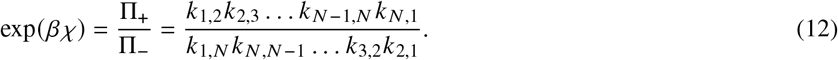

The rate constants in Equation 12 are treated as either first or pseudo first-order. A pseudo first-order reaction is a second order reaction for a molecule *A* that may exist in two states *i* and *j, A*_*i*_ +X → *A*_*j*_, where one reactant’s concentration [X] is much larger than the concentration of the other reactant and therefore treated as constant (35). In this limit, the concentration [X] is absorbed into the rate constant to form the concentration-dependent rate constant for a pseudo first-order reaction,

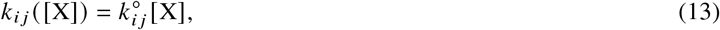

where 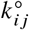 is the intrinsic rate constant, which is independent of concentrations. The rate constants may depend on other external thermodynamic forces, such as mechanical forces or forces imposed by a membrane potential but care has to be taken to formulate rate constant expressions in such a way that the external influence vanishes under equilibrium conditions. For simplicity, we include in the following only processes dependent on concentration but extension to other external driving forces is straightforward, as we will show in Results where we assess the effect of the membrane potential on active transmembrane transport. We expand the right-hand side of Equation 12 in terms of the intrinsic rates

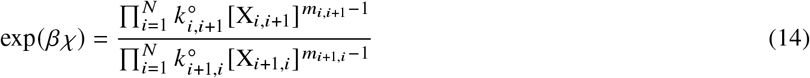

where X_*i,i*+1_ is the identity of the excess reactant in the reaction going from state *i* to *i* + 1. It is implied that indices exceeding *N* are cyclic (i.e., *N + i* is mapped to *i*). The reaction order *m*_*i j*_ for the transition from state *i* to state *j* is two for pseudo first-order reactions (e.g., binding of *X* to *A, A*_*i*_ +*X*_*i*_ →*A*_*j*_) and one for a first-order reactions (e.g., conformational change of *A, A*_*i*_ → *A*_*j*_). The driving potential separates into an intrinsic term, which only contains intrinsic rates 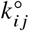, and an extrinsic term, which contains all the external variables such as the concentrations forming chemical gradients,

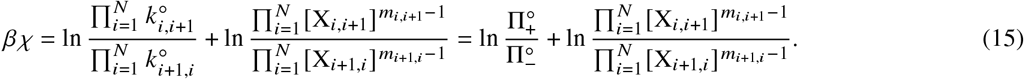

Without a chemical gradient and in the absence of any other driving forces the driving potential vanishes, and we are left with the equilibrium condition, Equation 1. Since the intrinsic rates do not depend on any of the extrinsic parameters and cycle driving forces of non-equilibrium systems must come from an external source of free energy, Equation 1 always applies to the intrinsic rates and thus constitutes a universal constraint on the 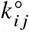, independent of the system being in equilibrium or out of equilibrium. By detailed balance, the logarithmic ratio of the intrinsic forward and backward rates equals the standard state free energy differences between the states, ln 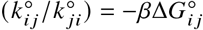; thus Equation 1 also expresses the constraint that the sum of the 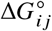 must vanish around any cycle (Equation 2). Overall, Equation 1 clearly shows that ratios of forward and backward rates of the cycle processes can not be determined independently (2); the corollary is that choosing a set of rates inconsistent with Equation 1 has the effect of introducing an additional non-physical cycle driving force (Equation 15) that will ultimately lead to incorrect predictions from the kinetic model.

Following Hill’s basic idea that we can use the equilibrium case to constrain the rates of a kinetic model we propose the following procedure to obtain thermodynamically consistent rates, given a set of input rates 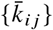 with errors 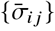 (e.g., obtained from experiments or simulations). These rates must describe the system in equilibrium. This condition is trivial to obey if the input rates are intrinsic second order rates as these can be converted to pseudo first order rates at standard state while ensuring that no external gradients are included. Note that the kinetic model should not contain any hidden states; in other words, all long-lived states must be included.

We want to determine a set of rates *k*_*ij*_ whose intrinsic components exactly obey Equation 1. We first calculate the free energy differences 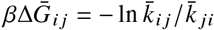 between all states to construct a potential graph using the maximum likelihood method in the previous section. To recover the rates from the free energies, we will project the inconsistent rates 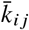 onto a set of consistent ones *k*_*ij*_ as shown next. Detailed balance requires that pairs of rates (*k*_*ij*_, *k*_*ji*_) must be consistent with

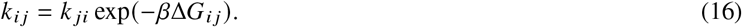

Since the consistency is only dependent on the ratio of forward and reverse rates, there exist infinitely many valid rate pairs. In order to select one rate pair we perform a maximum likelihood estimate for the model parameters {*k*_*ij*_, *k*_*ji*_} under the constraint Equation 16. We assume that the rate pairs have an underlying probability distribution that remains to be specified. We can then maximize the likelihood 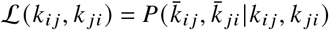, i.e., the probability to have observed the input rates given the model parameters {*k*_*ij*_, *k*_*ji*_} (and others as necessary) to obtain a single point estimate for the parameters of the distribution, the desired thermodynamically consistent rates *k*_*i j*_ and *k* _*ji*_.

We obtained the thermodynamically consistent Δ*G*_*ij*_ from another likelihood maximization (Equation 9) under the assumption that the free energy differences are normally distributed, Δ*G*_*ij*_ ∼ Normal (*μ, σ*) (considering Δ*G*_*i j*_ as a random variable in a temporary abuse of notation). If we rewrite Equation 16 as *K* = *k*_*ij*_ / *k*_*ji*_ = exp (−*β*Δ*G*_*ij*_) then *K* is a random variable that must be distributed according to the lognormal distribution. Given that the lognormal random variable *K* is a quotient of the random variables *k*_*ij*_ and *k*_*ji*_ it also follows that both *k*_*ij*_ and *k*_*ji*_ are lognormal. Thus, to be consistent with the assumptions that lead us to the free energy differences, the rates should be lognormal-distributed. Unfortunately, this assumption leads to unwieldy expressions (not shown) so we make the simplifying assumption that the rates are also normal-distributed, *k*_*ij*_ ∼ Normal(*μ*_*ij*_, *σ*_*ij*_). In general, the lognormal and the normal distribution behave rather differently (e.g., the support of the lognormal distribution is strictly positive as required for rates). However, for small variances relative to the mean 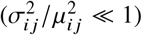 while in the vicinity of the mean, the leading term in a series expansion has the shape of a Gaussian with the same mean and standard deviation. Therefore, as long as the rate pair is already close to fulfilling the constraint (Equation 16) (i.e., the data being close to the mean of the distribution) we expect the simplifying assumption to be reasonable. To find the rates we will use maximum likelihood estimation with Equation 16 as an added constraint. Assume that we have *N*_*ij*_ samples *k*_*ij,n*_ of the rate of the transition *i* → *j* (and similarly *N*_*ji*_ for the reverse transition); we therefore also know the sample mean 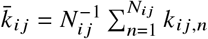 and variance while in the vicinity of the mean, the leading term in a series expansion has the shape of a Gaussian with the same mean and standard deviation. Therefore, as long as the rate pair is already close to fulfilling the constraint (Equation 16) (i.e., the data being close to the mean of the distribution) we expect the simplifying assumption to be reasonable. To find the rates we will use maximum likelihood estimation with Equation 16 as an added constraint. Assume that we have *N*_*ij*_ samples *k*_*ij,n*_ of the rate of the transition *i* → *j* (and similarly *N*_*ji*_ for the reverse transition); we therefore also know the sample mean 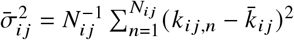. Under the assumption of the rates being normal-distributed, the likelihood reads

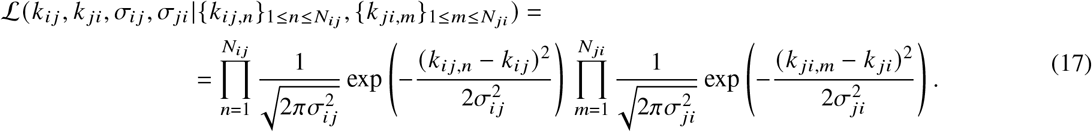

We then maximize the log-likelihood ln ℒ + *λ*(*k*_*ij*_ − *k*_*ji*_*K*) with the constraint added as a Lagrange multiplier to yield point estimates for the thermodynamically consistent rates

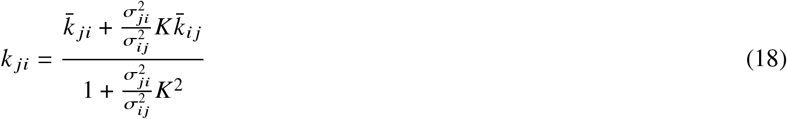

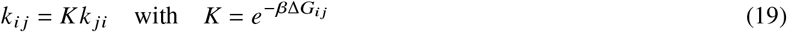

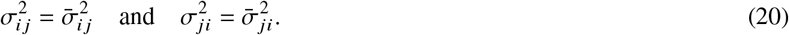

Note that the thermodynamically consistent parameters *k*_*i j*_, *k*_*ji*_, *σ*_*ij*_, *σ*_*ji*_ can be completely expressed in terms of the summary statistics of the rates, i.e., the 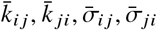, thanks to the assumption of being normal-distributed quantities. In a typical computational multiscale model approach it is common to have a single input value 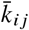 (with error estimate 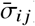) for a rate instead of a large number of rate measurements {*k*_*ij,n*_ }_1≤*n*≤*Nij*_, owing to the difficulty to obtain rate estimates in the first place. It is therefore especially convenient to be able to express the thermodynamically consistent rates in terms of single input rate estimates. Regardless of the exact underlying distribution, the rates in Equations 18 and 19 will always fulfill the constraint Equation 16 by construction.

The maximization procedure has a direct geometric interpretation, thanks to the symmetric shape of the underlying multivariate normal distribution. When viewed in the space of all possible rate pairs, a thermodynamically consistent rate pair must fall on the line through the origin with slope *K* = exp(−*β*Δ*G*_*ij*_), the solid black lines in figure 2. The maximum likelihood procedure minimizes the distance between the input rates and the consistency line in such a way that the input rates are projected along a line that passes through the point where a parallel of the consistency line forms a tangent to the error ellipse. This construction visualizes how smaller relative errors on the input rates restrain the choice of thermodynamically consistent rates more strongly. Consequently, rates with larger relative errors (indicated by longer axes in the Gaussian error ellipse) have more freedom to move towards consistency as shown in Figure 2.

**Figure 2:**
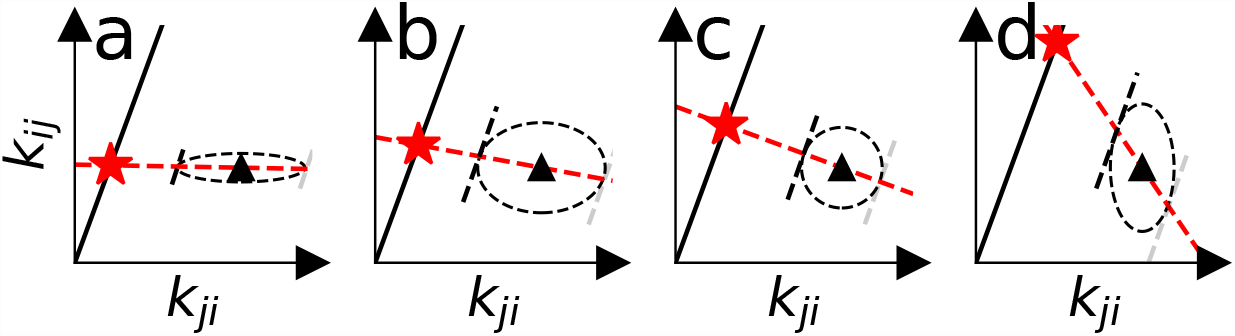
Geometric interpretation of rate likelihood maximization as a projection of input rate pairs (triangles) onto a thermodynamically consistent line (black line). The rate errors (depicted as the dashed ellipses) determine the final rate pair (star). **a** and **b** show higher certainty in the value of *k*_*i j*_ and show less movement of *k*_*i j*_ relative to *k* _*ji*_. **c** shows the case for equal certainty in both rates. **d** shows higher certainty in *k* _*ji*_.

By construction, the thermodynamically consistent rates *k*_*i j*_ exactly reproduce the free energy differences between states (Equation 16). The free energy differences were obtained via the potential graph approach (since the rates were required to describe an equilibrium system) and so they satisfy the cycle closure condition Equation 2. The intrinsic rates can be recovered from the thermodynamically consistent rates via Equation 13 if necessary. These intrinsic rates satisfy the kinetic cycle closure condition Equation 1 because in equilibrium it is equivalent to Equation 2, which they already satisfy. Thus, the *multibind* rate “projection” approach takes an initial set of rates, whose intrinsic components may violate the cycle closure condition Equation 1, and generates a set of rates that are thermodynamically consistent, which then can be used to analyze the kinetic graph under non-equilibrium conditions. The consistent rates contain information from all data and take errors of the input data into account via the global maximum likelihood approach and the rate projection step.

While the rate projection procedure guarantees thermodynamically consistent rates for a set of specified transitions, the model and its predictions of a physical system is sensitive to the omission of relevant states. During model construction, states may be excluded deliberately due to their short-lived nature and low impact on state probabilities or accidentally by virtue of limited system knowledge (e.g. a state may be hidden experimentally). In the case where the rates going from the known states to the missing state are equal, the prediction of the free energy *difference* between the major states, and thus their *relative* probabilities, are always correct, regardless of magnitude of the rates. If these rates are small, then the intermediate state is short-lived and does not affect state probability predictions. On the other hand, if these rates are large, then the missing state will have a substantial free energy contribution to the partition function and will affect the predictions of all state probabilities.

A full example of this scenario using mean first arrival times (36) to predict free energy differences and their deviations from the true free energy differences is developed in the Supplementary Information. In summary, in order to obtain correct thermodynamically consistent kinetic models, all major states and transitions must be included in the model: The *multibind* procedure cannot infer missing transitions or correct for missing information when generating thermodynamically consistent rates.

### Microscopic descriptors from macroscopic observables

The potential graph formulation for solving thermodynamic consistency comes with the advantage that the free energy differences defining the graph edges can easily become functions of an arbitrary number of external parameters such as pH and other ion concentrations. For the elementary binding reaction of a ligand X to a receptor A, A + X ⇌ A:X, the law of mass action relates the concentrations (or populations) of the free (dissociated) ligand and receptor ([X], [A]) to that of the associated (bound) state ([A:X])

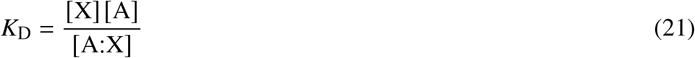

where the disassociation constant *K*_D_ reports the relative populations between the unbound and bound state and is independent of the concentrations. The free energy difference for the binding reaction, Δ*G*_bind_ = *G*_A:X_ − *G*_A+X_ = − *kT* ln [A:X] / [A], depends on the ligand concentration

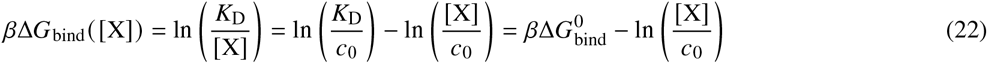

but can be written as the sum of the standard state free energy 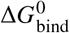 (relative to the standard state concentration *c*_0_, typically chosen as 1 M) with the concentration-dependent term. Equation 22 is general but in the special case of acid-base equilibria A^−^ + H^+^ ⇌ HA, i.e., proton binding, it is commonly expressed as

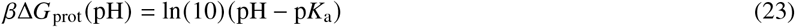

where the proton concentration is written as pH = −log_10_ [H^+^] / *c*_0_ and p*K*_a_ = −log_10_ *K*_*D*_ /*c*_0_ is the base-ten logarithm of the equilibrium constant.

Equations 22 and 23 describe the free energy difference between an occupied and an unoccupied binding site as a function of the binding ligand concentration. Macroscopic free energy differences are computed through the grouping of microstates describing a broader state definition (23, 24, 37, 38). The macroscopic free energy difference then takes the form

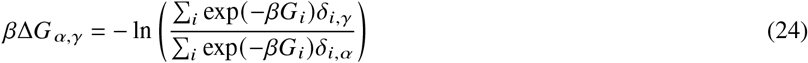

where *δ*_*i,α*_ and *δ*_*i,γ*_ are one (zero) if *i* is (is not) a member of the macrostate *α* or *γ* respectively. The *G*_*i*_ depend on concentrations/pH (and possibly other external parameters such as membrane potential) and so allow the calculations of the dependence of Δ*G*_*α,γ*_ on these external quantities. For example, if the thermodynamic cycle includes both protonation reactions and ligand binding reactions, the calculated ligand binding free energy Δ*G*_bind_ ([X]), pH will depend both on the ligand concentration and the pH.

### Inverse approach to obtaining microscopic quantities from macroscopic measurements

The consistent thermodynamic or kinetic microscopic *multibind* model can also be used in an inverse approach to infer microscopic quantities from macroscopic data and thus generate a microscopic model that is maximally consistent with measured data. The macroscopic reference (or training) data may come from experiments or simulations. In our inverse approach we use the *multibind* model to compute the macroscopic quantities from the microscopic descriptors, namely the free energy differences or the rates and then compare the computed quantities to the training data. The microscopic descriptors are then iteratively improved until the output of the model matches the training data to a prescribed tolerance.

As a proof of concept, we implement a simple Monte Carlo (MC) algorithm that minimizes the difference between the given and computed macroscopic features (Algorithm 1). For simplicity, we focus on microscopic free energies in an equilibrium thermodynamic graph but the generalization to a non-equilibrium kinetic model is straightforward. We initialize the potential graph with a set of free energy differences and generate the thermodynamically consistent model using the *multibind* approach. From the model, we calculate the macroscopic quantity of interest and determine the root mean squared difference (RMSD) to the training data. As long as the RMSD, which is used as a measure of goodness of fit, is above a set tolerance threshold *ϵ*, we modify the microscopic parameters of the model to generate a potentially better models. Here we change each microscopic parameter with a uniformly distributed random number *δ* from the symmetric interval –Δ ≤ *δ* ≤ Δ where Δ is chosen to generate sufficiently large changes. All edges in the graph are changed collectively because any single change will typically make the model thermodynamically inconsistent so it is more efficient to make one large collective set of changes instead of many smaller individual ones. If the model contains symmetries (e.g., due to symmetrical chemical environments) then sets of symmetry-related parameters are modified together with the same *δ*. These symmetries need to be accounted for by organizing parameters in lists of equivalent microscopic parameters (`symmetricEdges` in Algorithm 1). Any parameter without symmetries is contained in its own equivalency list. Based on the randomly modified parameters, the new thermodynamically consistent model is generated and the RMSD is recomputed. We use a zero-temperature Metropolis acceptance criterion whereby the current changes to the microscopic parameters are retained if the RMSD decreased compared to the RMSD of the last iteration. For the purposes shown in this work, such a criterion was sufficient to obtain a good match to experimental data. However, finite temperature MC can be easily implemented to generate a broader range of solutions. This procedure is repeated until the RMSD has decreased below the convergence threshold *ϵ* (or a maximum number of steps have been attempted without success).

#### Algorithm 1

Inverse *multibind* inference Monte Carlo algorithm. Microstates are represented as nodes in the potential graph and one microscopic model parameter is associated with each edge in the graph. `calculateRMSD()` calculates the root mean square deviation between the training data and the output of the model with the current set of microscopic parameters. The convergence tolerance for the iterative procedure is *ϵ*. Any edges that describe symmetrical processes (as determined by the user and listed in *symmetrical Edges*) are changed in the same manner, thus imposing symmetry on the solution. `Uniform` (− Δ, Δ) produces a uniformly distributed random number –Δ ≤ *δ* ≤ Δ. `solveMultibindModel()` implements the *multibind* procedure using Equation 9.

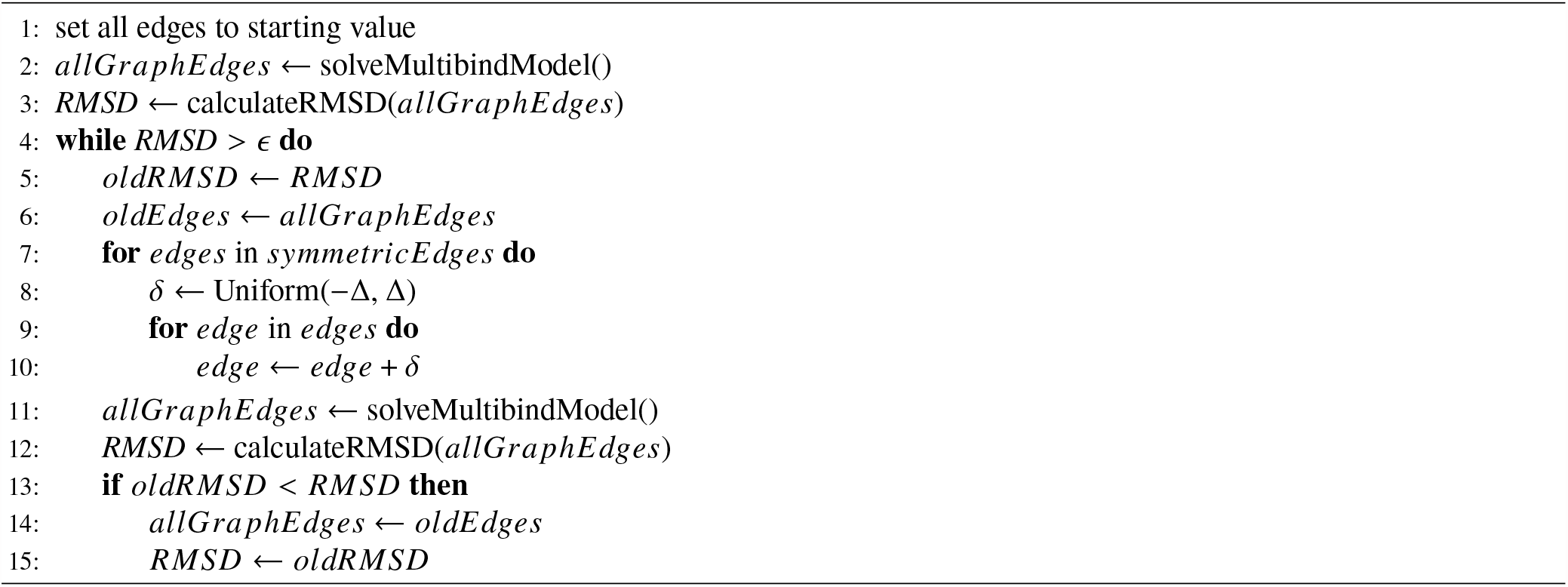

### Software implementation and Data Availability

The potential graph method and rate projection has been implemented in the multibind Python package, with source code available at https://github.com/Becksteinlab/multibind (code archived in the zenodo repository under DOI 10.5281/zenodo.5706366). Tests are implemented with PyTest and cover 100% of the code base. It is open source and released under the MIT license. It is built upon numpy (39), scipy (40), pandas (41), xarray (42), and networkx (43). Versions of Python supported include 3.8 and newer. With free energies defined at standard state concentrations, the model can be evaluated at many points in concentration space. States and their connecting processes are defined in a state and graph CSV file, respectively. `Multibind` objects, which are constructed from these two files, calculate and store free energies and the associated errors for the individual states. Proper labeling of macrostates allows for quick extraction of macrostate free energy differences from the resulting data structure. A wrapper class, `MultibindScanner`, assists in calculating state free energies over large concentration ranges and supports writing to the `xarray` NetCDF file format. Data used in this publication can be found in a separate repository at https://github.com/Becksteinlab/multibind-publication-code (archived in the zenodo repository under DOI 10.5281/zenodo.6927729).

## RESULTS/DISCUSSION

We will demonstrate the *multibind* approach for equilibrium and out-of-equilibrium systems. We begin with a toy model for a binding site with two protons in equilibrium in order to show how cooperativity, independence, and anti-cooperativity are captured by potential graph models. We demonstrate the inverse MC inference approach by calculating microscopic p*K*_a_s for a small molecule from experimental 1D NMR data. A simple model for a sodium/proton antiporter secondary active transport protein shows our approach to multiscale modeling in equilibrium. We then extend the antiporter model to out-of-equilibrium by considering driving forces due chemical and electrical potential differences and compute transport rates as function of the driving forces.

### Collective effects and cooperativity in a two-site system

We consider a simple toy system containing two proton binding sites (Fig. 3a), effectively a diprotic acid. In this model, the occupancy or vacancy of one binding site can potentially affect the binding of a proton to the other site. Therefore, four states (zero protons bound, proton bound to site 1, proton bound to site 2, both sites occupied by a proton) and four free energy differences between the four states (the microscopic p*K*_a_s) are needed to define the potential graph. Three sets of thermodynamically consistent p*K*_a_s (standard state free energy differences) were used as parameters in this model to show the coupling between binding sites (Figure 3).

**Figure 3:**
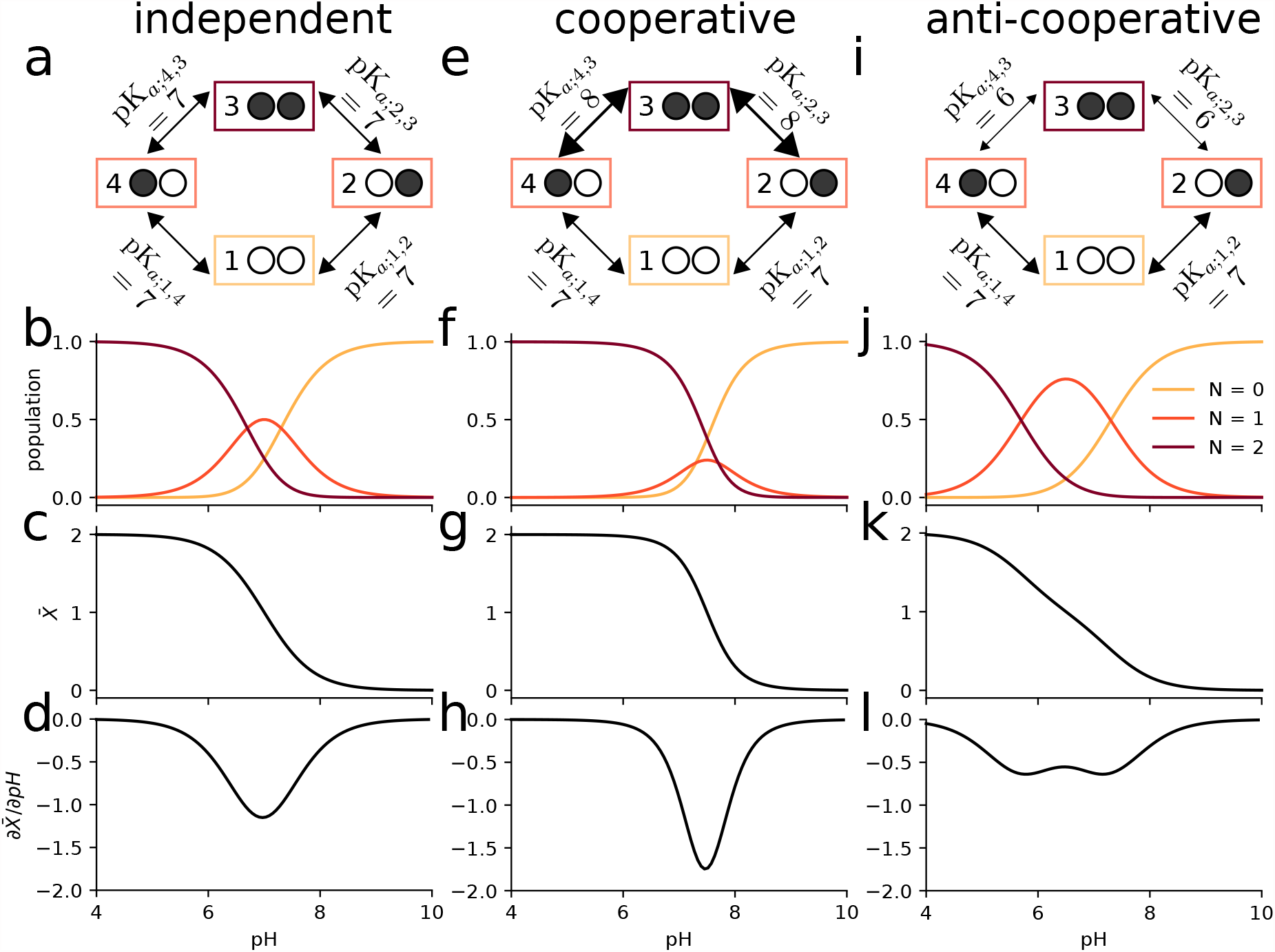
Two-site equilibrium model of proton binding. **a** Potential graph for *independent binding* with input p*K*_a_ values and microstates (numbered boxes); an occupied binding site is symbolized by a filled black circle whereas an unoccupied site is shown as an empty white circle. **b** Macrostate populations by number of protons (*N*) bound as function of pH for independent binding. **c** Mean proton occupancy 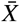 of the system as a function of pH for independent binding. **d** Rate of uptake 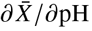 as function of pH for independent binding. **e** Potential graph for *cooperative binding*. **f** Macrostate populations for cooperative binding. **g** Mean proton occupancy for cooperative binding. **h** Rate of proton uptake for cooperative binding. **i** Potential graph for *anti-cooperative binding*. **j** Macrostate populations for anti-cooperative binding. **k** Mean proton occupancy for anti-cooperative binding. **l** Rate of proton uptake for anti-cooperative binding.

In the first case, where the binding of a proton to one site has no effect on the binding free energy of the other site, all p*K*_a_ values were set to 7 (Figure 3a). The pH dependent free energies for each state are determined through the maximum likelihood estimator (Equation 9), which in this case preserved the input pH dependent free energy differences (Equation 23) since they already satisfied cycle closure. Macrostate populations for the proton bound states (total number of protons *N* = 0, *N* = 1, or *N* = 2), were calculated using Equation 7 (Figure 3b). Starting from a fully occupied system at pH = 4, increasing the pH promotes the less occupied *N* < 2 states. Once the pH is sufficiently large (> 9), the observed system will almost always be entirely deprotonated. The weighted sum of the macrostate probabilities was calculated to predict the mean proton occupancy 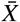 of the system as a function of pH (Figure 3c) along with the “rate of uptake” with respect to a changing pH, 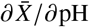 (Figure 3d), which characterizes the strength of response of the binding site to changes in pH and can be used to quantify cooperativity (38). The example of independent binding establishes a baseline for the behavior of this simple model when considering cooperative and anti-cooperative binding next.

We then modeled a system where the p*K*_a_ of one site was dependent on the state of the other site, namely binding in one site could either increase or decrease binding to the other site. For simplicity, we treated the two sites as symmetrical. For the cooperative model, binding of the first proton upshifts the p*K*_a_ for the second proton binding event (Figure 3e), i.e., it lowers the free energy of binding (Equation 23) and so increases the probability of binding a second proton when a first one is already bound. When the p*K*_a_ is instead downshifted (Figure 3i), the binding free energy increases, binding of the second protein becomes less likely, and anti-cooperative behavior emerges, as becomes clear when analyzing state probabilities.

The probabilities to observe a specific microstate (Equation 6) provide a full microscopic picture of the system as a function of pH. The microstates are aggregated by the total number of protons bound *N* to produce the corresponding macrostate probabilities (Equation 7). In the cooperative case, the intermediate *N* = 1 proton state is suppressed (Figure 3f) compared to the case of independent binding (Figure 3b). Cooperativity increases the slope of the deprotonation curve (Figure 3g), which is seen directly as a stronger (more negative) rate of uptake (Figure 3h).

In the anti-cooperative case, the single bound macrostate is promoted compared to independent binding and becomes dominant over a wider pH range (Figure 3j); the deprotonation curve becomes flatter and starts exhibiting a double-step shape (Figure 3k) and the proton rate of uptake decreases in magnitude (Figures 3l).

This simple example demonstrates that the potential graph formalism can capture cooperative or anti-cooperative behavior. It clearly shows that such behavior is a direct consequence of microscopic (standard state) free energy differences 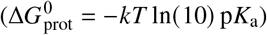 or, equivalently, equilibrium constants 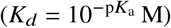. The model is also sufficiently simple to demonstrate by inspection that the choice of p*K*_a_ values cannot be arbitrary and only three out of the four p*K*_a_ values can be chosen. The fourth one must be chosen to satisfy the cycle closure property, which in this case reads (using Equation 23) Δ*G*_14_ + Δ*G*_43_− Δ*G*_23_ −Δ*G*_12_ = −*kT* ln (10) (p*K*_a;14_ + p*K*_a;43_ − p*K*_a;23_ − p*K*_a;12_) = 0. Although the cycle closure requirement is obvious here, it becomes more complicated to satisfy in models with multiple cycles. In all these cases, the *multibind* approach will ensure a thermodynamically consistent model via the potential graph.

### Extracting microscopic p*K* _a_ values from experimental 1D NMR data

A more complicated model for proton binding is necessary for the organic molecule diethylenetriaminepentaacetic acid (DTPA) (Figure 4a). Protonation of the three amine nitrogen atoms was measured by 1D NMR (44). The experiment distinguished binding to the central nitrogen from binding to the two symmetrical terminal nitrogens. The site-specific protonation curves as function of pH (reproduced in Figure 4b, c) show distinct non-monotonic behavior for the central nitrogen and clear deviation from the simple independent binding behavior seen in the simple two-site model (Figure 3c), thus indicating coupling between the sites. Previous work showed that microscopic p*K*_a_ values could be extracted from the NMR data using a “decoupled site representation” (45). Here we demonstrate that we can use the *inverse multibind* approach to directly infer the microscopic p*K*_a_ values from the 1D NMR data.

**Figure 4:**
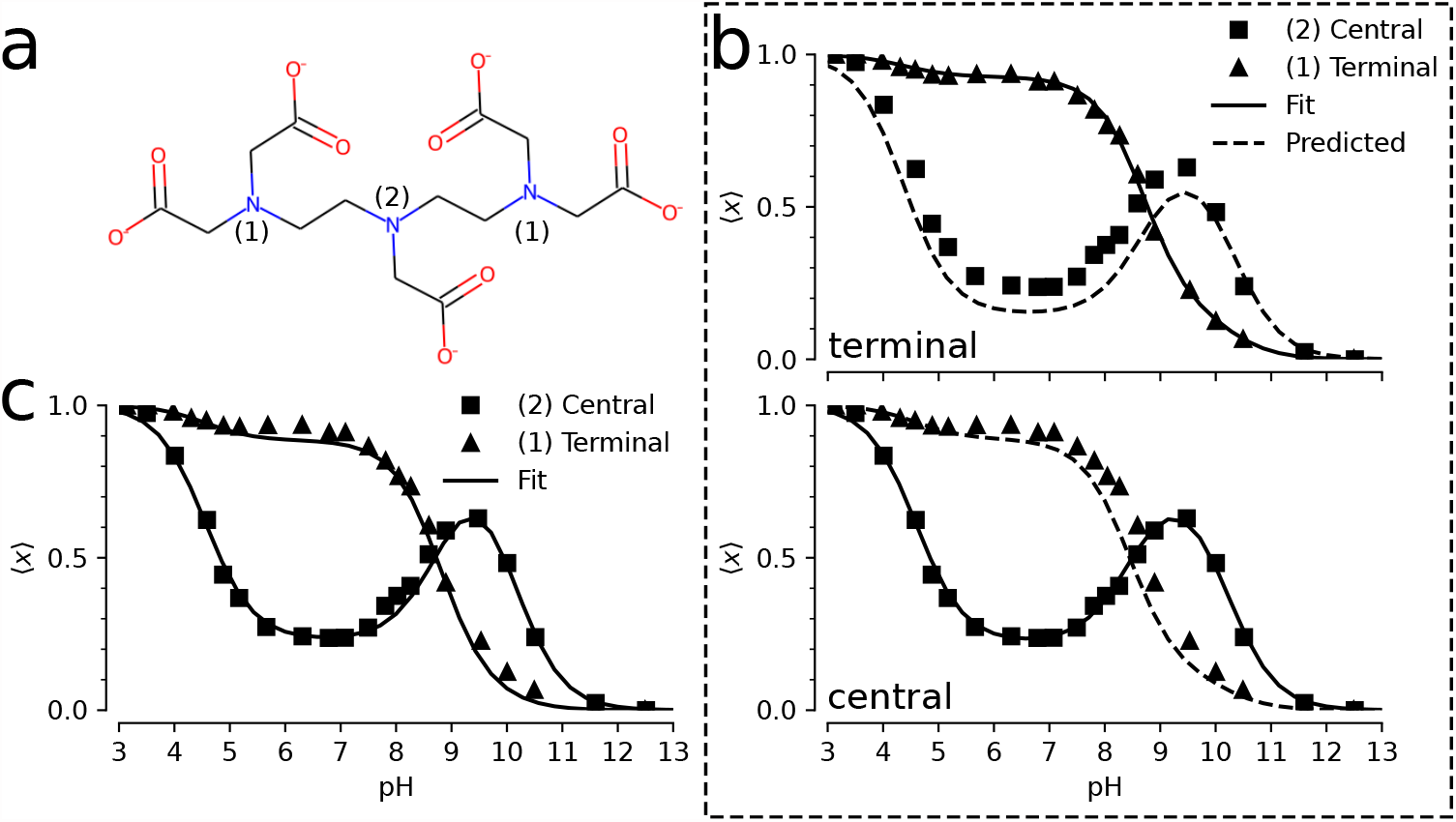
*Inverse multibind* approach for inferring microscopic p*K*_a_ values from 1D NMR data for DTPA.**a** Chemical structure of diethylenetriaminepentaacetic acid (DTPA) with protonation sites at terminal (1) and central (2) nitrogen atoms indicated. **b** Mean proton occupancy of protonation sites as function of pH. Experimental 1D NMR data from Submeier and Reilley (44) are shown as squares for the central site and triangles for the symmetrical terminal sites. Black solid lines are the *inverse multibind* fits to the data. Black dashed lines are predicted occupancy curves from the model. Top panel: only experimental data for terminal nitrogens (triangles) were used to generate the microscopic model. Bottom panel: only experimental data for the central nitrogen were used. **c** Mean proton occupancy (as b) when experimental data from all sites were used to infer the microscopic model.

We describe the DTPA system as a 3-site model with 2^3^ = 8 micro states (Figure 5a) as an extension of the 2-site model shown in the previous section. States were labeled in binary notation, where three sites are represented as a three-digit binary number with a proton-occupied site set to 1 and an empty site set to 0; for instance, a proton bound to the central site is state 010. We applied a simple zero-temperature Metropolis-Hastings Monte Carlo algorithm to sample p*K*_a_ parameter space for a range of pH values, as described in more details in Methods. We initialized the model with the identical p*K*_a_=7 for all microscopic protonation reactions. Given a set of p*K*_a_ values, the computed titration curves were compared to the experimental curves, which were manually digitized from Submeier and Reilley (44) with the WebPlotDigitizer tool (46) with all data included in the Supplemental Table S1. A new set of p*K*_a_ values were chosen at each step and accepted if the root-mean-square deviation to the experimental data decreased. In the MC trial step, p*K*_a_ shifts were chosen uniformly between ± 2 pH units (Δ = 2). Due to the symmetry of the terminal nitrogen atoms, the p*K*_a_s involving the terminal nitrogen sites were updated in the same manner to maintain the symmetry. Minimization was performed to a final RMSD of 0.029. This process yielded a set of microscopic p*K*_a_s (Table 1) that fit the experimental data closely (Figure 4c). In particular, the non-monotonic behavior of the central site was captured accurately.

**Table 1:** Microscopic p*K*_a_ values inferred from all DTPA NMR data with the *inverse multibind* MC approach. The subscript and superscript denote the starting and ending states, respectively. p*K*_a_ values are grouped by how the total number of protons bound to the three nitrogens changes.

| $0 \rightarrow 1$ | $1 \rightarrow 2$ | $2 \rightarrow 3$ |
| --- | --- | --- |
| $pK_{a_{000}}^{001} = 9.0$ | $pK_{a_{001}}^{011} = 9.0$ | $pK_{a_{011}}^{111} = 5.5$ |
| $pK_{a_{000}}^{010} = 10.1$ | $pK_{a_{001}}^{101} = 9.8$ | $pK_{a_{101}}^{111} = 4.7$ |
| $pK_{a_{000}}^{100} = 9.0$ | $pK_{a_{010}}^{011} = 7.9$ | $pK_{a_{110}}^{111} = 5.5$ |
| | $pK_{a_{010}}^{110} = 7.9$ | |
| | $pK_{a_{100}}^{101} = 9.8$ | |
| | $pK_{a_{100}}^{110} = 9.0$ | |

**Figure 5:**
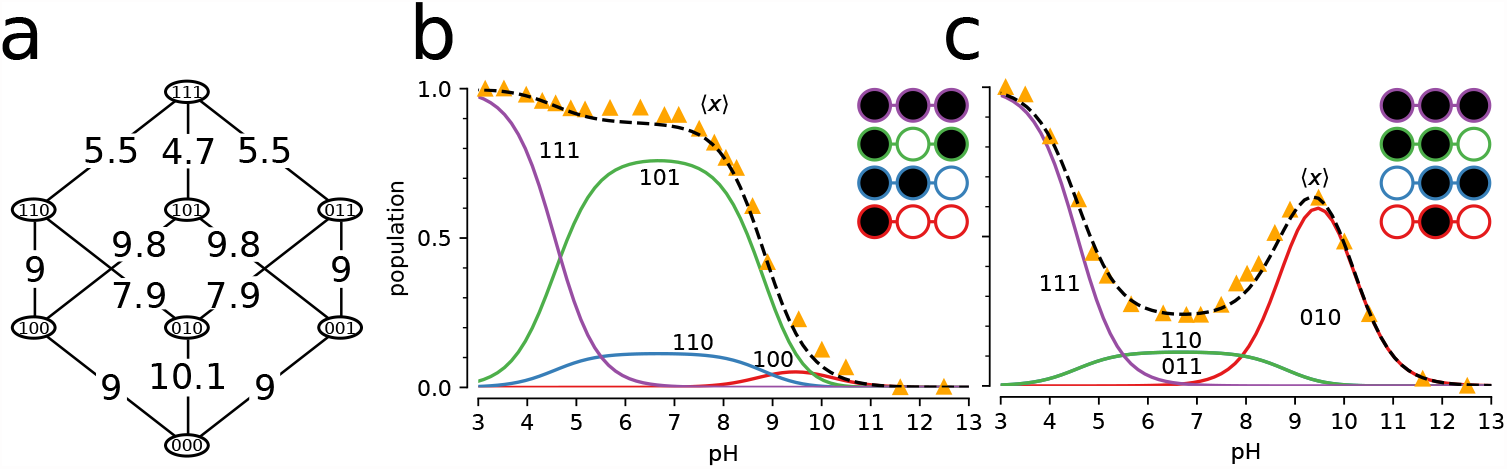
Microscopic model for protonation of DTPA. **a** Microscopic potential graph with p*K*_a_ values inferred from all experimental 1D NMR data. States are labeled in binary notation where the sites (terminal/central/terminal) correspond to the digits and a proton-bound site is denoted by a 1 and an unoccupied one by 0. **b** Microstate populations with at least one terminal nitrogen protonated as function of pH. The microstates are color coded and shown as filled (protonated) and empty (deprotonated) circles in the inset. The experimental ensemble titration curves are shown as orange triangles and the predicted curves as dashed lines. **c** Microstate populations with the central nitrogen protonated.

We evaluated the robustness of the inference by asking if a reasonable set of p*K*_a_s still be inferred even if not all site populations are directly measured. We performed the same MC inference on a subset of the data on used only one of the two experimental titration curves as training data while the other one was used for validation. As shown in Figure 4b, even in the case of limited training data, the model is able to predict the shape of the validation titration curve. Even when the apparently featureless titration curve for the terminal nitrogens is used for training, the model predicts the unusual shape of the central nitrogen curve (Figure 4b, top panel), indicating that each curve contains sufficient information about the cooperativity of the proton binding sites. The inferred microscopic p*K*_a_s and their fitting RMSDs are tabulated in Supplemental Tables S2 and S3 of the supporting information, respectively.

Apart from recovering the macroscopic features of the system, the MC inference enables interpretation of microscopic descriptors and their coupling. For instance, the nonstandard shape of the titration curve for the central nitrogen is explained by suppression of the two doubly protonated titration states with a proton bound to the central nitrogen (Figure 5b, c). For the *N* = 2 macrostate, the system is most likely found in the doubly protonated terminal nitrogen state. The p*K*_a_ for the central nitrogen, in the absence of either terminal nitrogen is 10.1 (Figure 5a and Table 1). In the event that either terminal nitrogen is protonated, this p*K*_a_ shifts to 9.0, in contrast to the initial p*K*_a_ of 9.0 for the terminal nitrogen atoms. One protonated terminal nitrogen will shift the p*K*_a_ of the other to 9.8. From these data, we conclude that the proton binding to a terminal nitrogen atom and the central nitrogen are anti-cooperative while the proton binding of the terminal nitrogen atoms are cooperative.

In summary, the *inverse multibind* approach can be used to infer microscopic properties of polyprotic acids from macroscopic measurements such as titration curves. However, the approach is not limited to acids and protonation events. Because multiple species can be described together in a consistent thermodynamic model, our inference approach has the potential to provide additional microscopic details in applications such as pH-dependent ligand binding or coupled ion/proton transport.

### Multi-ligand model and kinetics for a secondary active antiporter

As an example for using *multibind* in a bottom-up multiscale approach we created a simplified generic model of a sodium/proton antiporter, a type of secondary active transporter that uses the free energy in the electrochemical sodium transmembrane gradient to drive the energetically unfavorable transport of protons in the opposite direction (47–49). Here our emphasis is on demonstrating general principles so instead of specific parameters, we use reasonable estimates for rates in a simplified single-cycle kinetic model (Figure 6a). Using the *multibind* approach ensures that the resulting model is thermodynamically and kinetically consistent, even though the estimates were not.

**Figure 6:**
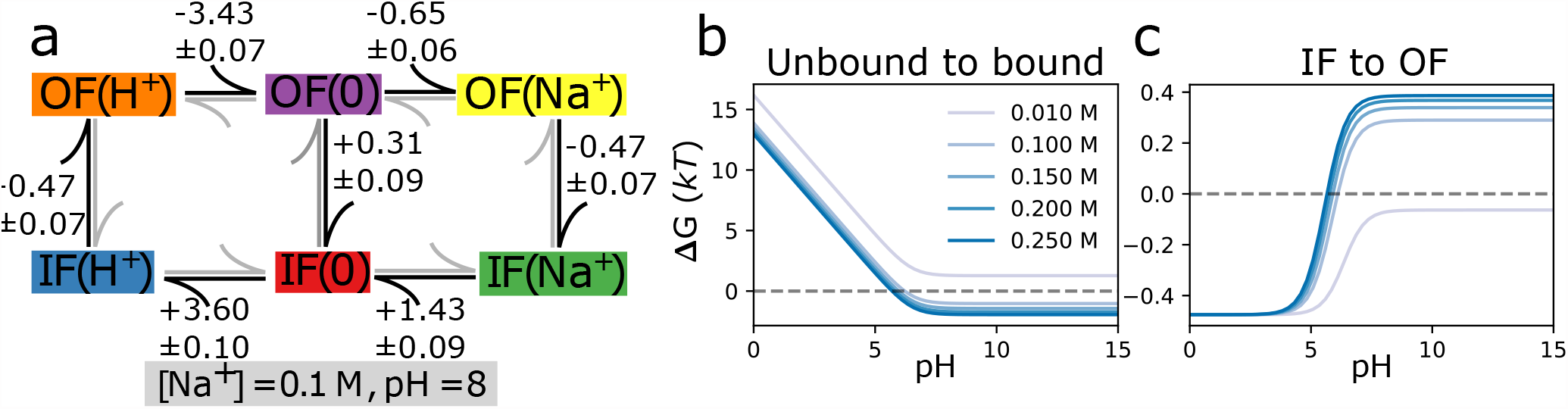
Macroscopic free energy predictions for a toy antiporter model with no concentration gradients or membrane potential. The sum of the differences (in units of *kT*) around the loop (**a**) vanish, and therefore the graph obeys detailed balance. These differences were the result of the *multibind* approach using starting values from Table 2 defined at a sodium concentration of 100 mM and pH 8, and whose sum did not vanish. Having solved for the state free energies, macroscopic features of the system are predicted. (**b**) shows the sodium binding free energy while (**c**) shows the inward to outward facing binding free energy difference as a function of the pH and sodium concentration.

Like all known sodium/proton antiporters, our toy antiporter functions according to the alternating access mechanism whereby a conformational transition between the outward facing (OF) and inward facing (IF) conformation exposes the binding site to either the extracellular or the intracellular solution (49). Furthermore, we assume that sodium ions and protons compete for the same binding site, a common principle in this specific family of transporters (48). Therefore, our model contains OF and IF states with either H^+^ or Na^+^ bound. Transitions between the IF and OF conformations are possible in any binding state, although the transition rates for conformational changes with an empty binding site (a leak edge) are much lower than those including bound ligands, modeling a strongly coupled antiporter. The coupling between the transported and driving ion has been addressed in previous works by Berlaga and Kolomeisky (10, 11), where chemical-kinetic models of antiporters and symporters were developed to analytically explore transport properties. This coupling was quantified by a dimensionless parameter, *x*, which is the ratio of the ion transporting conformational change rates and the leaking conformational change rates. In the present work, *x* falls approximately between 50 and 80 for a small leak at equilibrium (and no membrane voltage) and is indicative of strong coupling. However, since our model contains the membrane voltage, which modifies the ion-bound conformational change rates, and does not employ many of the simplifying assumptions of Berlaga and Kolomeisky (11), which made their model analytically tractable, a direct comparison of the coupling strength is difficult.

We initially consider only electroneutral transport, according to the outer cycle, where one proton is exchanged for one sodium ion (Figure 6a). With the ultimate goal of determining transport rates in non-equilibrium conditions, the primary set of inputs are forward and reverse rates, which can later be related to p*K*_a_s and *K*_D_ through the free energy and detailed balance for intermediate steps. Rates involving ligand binding are treated as pseudo first-order rates and are linearly dependent on the ligand concentration. Proton or ion release (“unbinding”) is treated as independent of concentration.

First, we defined the order of magnitude forward and backward rates of each process along with their errors, tabulated in Table 2, and reflect realistic pH (8) and sodium concentrations (100 mM). The inward and outward binding site p*K*_a_s were set to 6.4 and 6.5, respectively, and fall within a reasonable p*K*_a_ range for modeling the binding site of a sodium proton antiporter (50–52). The sodium binding constant *K*_*D*_ for the inward and outward facing states are, respectively, 24 mM and 52 mM, in line with typical *K*_*D*_ value ranging from 1.6 mM to 30 mM (52, 53). The on-rates for proton and sodium binding were estimated assuming diffusion limited binding by solving the steady-state Smoluchowski equation for a disk-shaped binding site as described in the Supplementary Information. The off-rate for unbinding is then calculated from the on-rate and the free energy difference assuming mass-action kinetics by Equation 16. Note that any thermodynamic inconsistencies in the binding free energies used are then reflected in the ratio of the rates. The conformational change rates were chosen to be within an order of magnitude of the sub-millisecond transport rates of secondary active transporters (49). The forward and reverse rates between adjacent states were converted into free energy differences, which were then made consistent through the application of the maximum-likelihood method and used to project the input rates onto a new set of thermodynamically consistent ones, tabulated in Table 2.

**Table 2:** First order rate constants for the sodium/proton antiporter model at [Na^+^]_in_ = [Na^+^]_out_ = 100 mM and pH_in_ = pH_out_ = 8. Input forward 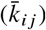 and backward 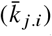 rates with estimated errors were determined as described in the text and corresponding free energy differences 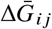 were calculated from these rates with error propagation. The thermodynamically consistent parameters Δ*G*_*ij*_, *k*_*ij*_, and *k*_*ji*_ were determined with the potential graph approach together with rate projection. The intrinsic rate constants 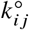 of the model are listed in Supplemental Table S5.

| State 1 | State 2 | input parameters |  |  |
| --- | --- | --- | --- | --- |
| | | $\bar{k}_{12} \pm \bar{\sigma}_{12} \text{ (s}^{-1}\text{)}$ | $\bar{k}_{21} \pm \bar{\sigma}_{21} \text{ (s}^{-1}\text{)}$ | $\Delta\bar{G}_{12} \pm \bar{\sigma}_{\Delta G} \text{ (kT)}$ |
| IF(H<sup>+</sup>) | OF(H<sup>+</sup>) | 8000.00 ± 100.00 | 5000.00 ± 100.00 | -0.470 ± 0.024 |
| OF(H<sup>+</sup>) | OF(0) | 6000.00 ± 100.00 | 200.00 ± 15.00 | -3.401 ± 0.077 |
| OF(0) | OF(Na<sup>+</sup>) | 320000000.00 ± 1700000.00 | 170000000.00 ± 10000000.00 | -0.633 ± 0.059 |
| OF(Na<sup>+</sup>) | IF(Na<sup>+</sup>) | 8000.00 ± 100.00 | 5000.00 ± 100.00 | -0.470 ± 0.024 |
| IF(Na<sup>+</sup>) | IF(0) | 70000000.00 ± 9500000.00 | 320000000.00 ± 1700000.00 | 1.520 ± 0.136 |
| IF(0) | IF(H<sup>+</sup>) | 200.00 ± 25.00 | 8000.00 ± 400.00 | 3.689 ± 0.135 |
| OF(0) | IF(0) | 100.00 ± 31.62 | 135.00 ± 31.62 | 0.300 ± 0.394 |
| thermodynamically consistent parameters |  |  |  |  |
| | | $k_{12} \pm \sigma_{12} \text{ (s}^{-1}\text{)}$ | $k_{21} \pm \sigma_{21} \text{ (s}^{-1}\text{)}$ | $\Delta G_{12} \pm \sigma_{\Delta G} \text{ (kT)}$ |
| IF(H<sup>+</sup>) | OF(H<sup>+</sup>) | 8006.21 ± 12.57 | 4990.04 ± 19.14 | -0.473 ± 0.071 |
| OF(H<sup>+</sup>) | OF(0) | 6007.96 ± 4.12 | 194.47 ± 3.57 | -3.431 ± 0.065 |
| OF(0) | OF(Na<sup>+</sup>) | 320044062.77 ± 392.95 | 167079476.75 ± 3240.18 | -0.650 ± 0.063 |
| OF(Na<sup>+</sup>) | IF(Na<sup>+</sup>) | 8006.24 ± 12.15 | 4989.98 ± 18.49 | -0.473 ± 0.066 |
| IF(Na<sup>+</sup>) | IF(0) | 76760846.63 ± 2645.76 | 319948058.80 ± 6426.80 | 1.427 ± 0.091 |
| IF(0) | IF(H<sup>+</sup>) | 215.85 ± 4.08 | 7889.02 ± 30.87 | 3.599 ± 0.093 |
| OF(0) | IF(0) | 99.70 ± 2.40 | 135.22 ± 3.79 | 0.305 ± 0.089 |

To study the model in equilibrium, the resulting free energy differences from the maximum-likelihood approach were converted into p*K*_a_s and *K*_D_s for the ligand binding transitions (Equations 23 and 22), which can be found in the Supplemental Table S4. In equilibrium, using just the thermodynamically consistent free energy differences (Figure 6a and Table 2), we assessed the influence of pH and sodium concentration on observing different states of the systems. The probability to observe a sodium-bound state, quantified through the sodium binding free energy Δ*G*, depends strongly on pH below around pH 7 but then remains constant (Figure 6b) because at low pH, protons compete with sodium ions for the binding site and thus at low pH, sodium binds only weakly. A decrease in sodium concentration leads to a decrease in binding free energy and at values below the lowest sodium *K*_*D*_ ≈ 24 mM (see Supplemental Table S4) no binding occurs, as expected. At high pH, too few protons are available to compete and the binding free energy becomes independent of the proton concentration. The conformational equilibrium between IF and OF (also quantified by the free energy Δ*G* between the two states) could be shifted from favoring the IF conformation at higher sodium concentrations to favoring OF at lower concentrations (Figure 6c). This result demonstrates, with a simple model, how different environmental conditions may bias a transporter to appear in different conformations, as sometimes observed in the variation of conformations with crystallization conditions in X-ray crystal structures.

In order to operate, secondary active transporters require non-equilibrium conditions. By establishing a membrane potential along with sodium and proton gradients, the model captured non-equilibrium behavior. Rates involving the movement of an ion across an electric potential were modified such that a kinetic barrier is placed halfway through the membrane and thus the rate took the following form (54),

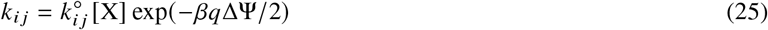

where *q* is the transported charge and ΔΨ = Ψ_in_− Ψ_out_ is the membrane potential.

In order to calculate the transporter turnover number from the model, we calculated the net transition flux between proton-bound OF and IF conformations by solving the underlying master equations under steady state conditions as described in Supplementary Information. Here positive values of the turnover number indicate proton transport out of the cell while negative values indicate proton transport into the cell. Note that for our simple antiporter model, the proton and sodium fluxes are equal only when there is perfect coupling.

The internal sodium concentration was set to 10 mM and the internal and external pH were set to 7.4 and 7, respectively. The external sodium concentration and the membrane potential were allowed to vary in order to mimic an experimental setup. Varying the external sodium concentration and the membrane potential (Fig. 7a, 7b), the steady state solutions of the kinetic model show that as the external sodium concentration is increased, positive turnover is increased and overcomes the drive resistance present in the pH gradient. This was calculated across a range of membrane voltages where a positive voltage indicates a lower potential on the inside of the cell. At large membrane potentials, transport is stalled due to the rates of conformational change approaching zero exponentially, since those processes involve moving charges across an electric potential. The direction of transport is entirely dependent on the pH and sodium concentration gradients in the electroneutral case as no net charge is moved across the membrane potential over the cycle. This is not to say that the membrane voltage does not assist in transport in this model. In fact, we saw that a positive membrane voltage amplified transport even when the sodium concentration gradient was removed at [Na^+^]_out_ = 10 mM.

**Figure 7:**
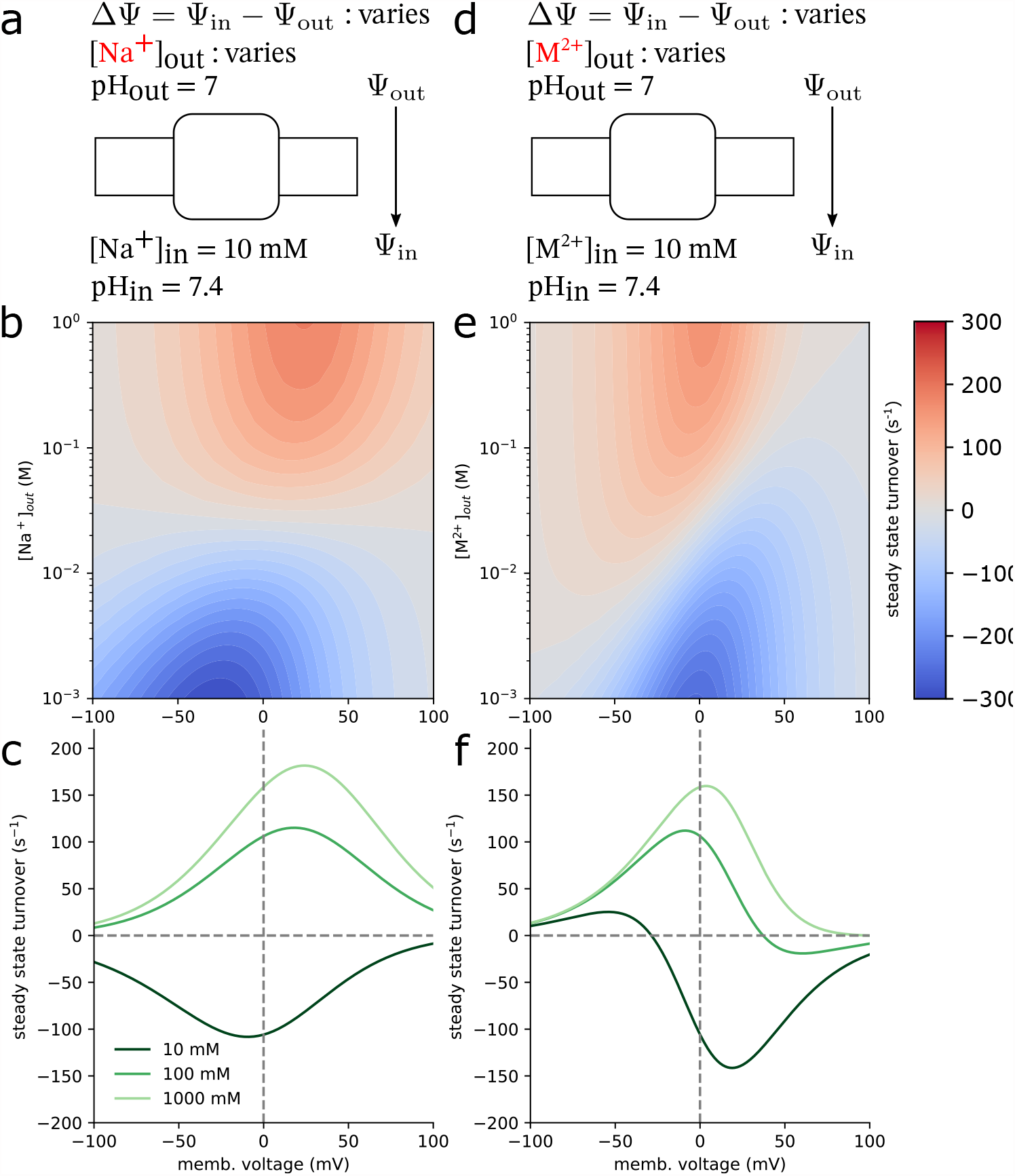
Antiporter turnover numbers (Na^+^/H^+^ or M^2+^/H^+^ transport cycle completions per unit time) for the electroneutral (Na^+^/H^+^, **a**) and electrogenic (M^2+^/H^+^, **d**) antiporter models in steady state as a function of the cycle driving force in the driving ion gradient and an applied membrane voltage. The internal *driving* ion concentration was fixed at 10 mM for both models while the external concentration was a model parameter ranging from 1 mM to 1 M. Internal and external pH values were respectively set to 7.4 and 7. The membrane voltage, ΔΨ = Ψ_in_− Ψ_out_, which ranged from -100 mV to 100 mV. Turnover numbers were calculated for both the electroneutral (**b**) and electrogenic (**e**) models as a function of the external driving ion concentration and the applied membrane voltage. Turnover numbers at fixed external driving ion concentrations are shown as a function of the membrane voltage for both models (**c** and **f**).

To demonstrate electrogenic transport, we changed the identity of the monovalent sodium ion from the electroneutral case to a divalent cation, M^2+^, thus modeling an electrogenic divalent cation/proton antiporter(55–57) (although the transport stoichiometries are not necessarily 1:1 in all cases). In order to highlight the effects due to a net charge transport, all of the intrinsic rates were kept the same from the previous electroneutral case and turnover numbers were again calculated (Figure 7e,7f). In the case where no metal cation concentration gradient exists ([M^2+^] _out_ = [M^2+^] _in_ = 10 mM) and the membrane potential is zero, protons will flow in reverse (Figure 7f, dark green line). Decreasing the membrane potential from this point begins to stall net transport as the turnover root is approached. The membrane potential relative to this point determines the direction of transport. Any positive change leads to reversed transport while any negative change leads to forward transport. Large membrane potentials in either direction stall transport. In the large negative case, the movement of metal cations from the exterior to the interior of the cell approaches zero. Large positive membrane potentials halt transport of protons from the interior to the exterior. The sign of the cycle drive, χ (Equation 12), determines the transport direction. In the electroneutral Na^+^:H^+^ case, the contribution due to the membrane potential vanishes from the ratio of forward and reverse rates due identical translated charges (Equation 25). In contrast, the cycle drive of the electrogenic M^2+^:H^+^ case is a linear function of the membrane potential and is proportional to the charge difference between the transported ions (Equation S13). This means that the electroneutral transporter’s outer cycle direction is determined entirely by the chemical gradients of the transported ions and cannot be reversed by applying a membrane potential. On the other hand, the electrogenic transporter’s outer cycle drive is a balance between the the chemical gradients and the electric term describing the movement of a net charge across the applied potential.

The overall transport stoichiometry in both cases is also a function of the leak cycle pathway between the empty inward and outward conformations. By varying the magnitude of the rates of leak pathway, using the values tabulated in Table S6, the antiporter’s electrogenicity also changes. When the leak pathway is removed, the proton transport numbers remain practically the same (Figure S2). When a large leak is introduced, with transitions occurring on the order of 1000 s^-1^, cycles bypassing the transport of ions across an unfavorable electric potential become significant and dramatically increases system complexity. Note that when a leak pathway is present, it is possible, on average, for both ions to be transported in the same direction, leading to competition for the limited state flux passing through the leak edge. In the electroneutral case, these effects are most apparent in the low sodium concentration (10 mM) and large negative membrane voltage regimes (−100 to -50 mV), but diminish with increasing sodium concentrations. Where reversed transport was once quickly stalled by unfavorably moving sodium out of the cell (albeit with highly coupled transport), against the membrane potential, transport is amplified since transitions across the leak pathway are unaffected by this potential (Figure S3a, S3b). The metal cation driving ion model shows similar behavior across the same concentration and voltage ranges. The observed reversal of the proton transport direction observed with a small leak disappears since the favorable movement of the M^2+^ ions no longer requires the unfavorable movement of protons out of the cell (Figure S3c, S3d).

## CONCLUSIONS

We introduce a framework where non-equilibrium kinetic networks can be constructed by first building underlying equilibrium potential graphs, emphasizing the states directly instead of the edges connecting them. By solving for thermodynamically consistent free energy differences with a maximum likelihood approach, all input uncertainties are used and path dependent state free energies are eliminated. The potential graph formulation includes chemical degrees of freedom that are typically hidden from experimental measurements but, nevertheless, lead to the macroscopic measurements that are accessible to experiments. Through a microscopic description, rather than a macroscopic description, the coupling between processes are revealed and may lead to insights related to molecular mechanisms.

We demonstrate that thermodynamically consistent kinetic rates can then be systematically determined by another maximum likelihood procedure that can be understood as projecting an initial set of thermodynamically inconsistent rates pairs onto a set of rate pairs that satisfy detailed balance. By building kinetic networks this way, artificial and incorrect driving forces, which would originate from rates that fail to obey detailed balance, are eliminated.

From an equilibrium and purely thermodynamic standpoint, we show that the potential graph formulation is capable of modeling cooperativity that can have sizable effects on macroscopic behaviors. A cooperative relationship between processes leads to suppression of intermediate state populations while an anti-cooperative relationship promotes intermediate states. These effects can directly impact concentration dependent macroscopic observables such as transport stoichiometry in transporter proteins. These models allow exploration of which degrees of freedom contribute to overall function by providing a microscopic model containing state populations and transition fluxes.

We demonstrate that predicted macroscopic observables can be used to infer a system’s microscopic descriptors when comparing those observables to experimental findings. We further show that a simple Metropolis-Hastings Monte-Carlo optimizer can construct a potential graph that continuously enforces a thermodynamic consistent model. We demonstrated this method by constructing a three site (eight state) model and determining a set of p*K*_a_s that minimized the root-mean-square deviation between predicted and titration curves from an NMR experiment. This application showed that these models are capable of recreating titration curves that deviate from ideal Henderson-Hasselbalch curves in a robust manner. The inverse *multibind* approach promises to be a useful tool to infer microscopic properties from experimental data whose degrees of freedom may be coupled in complex and subtle ways.

Using a simplified sodium/proton antiporter model we showed how a set of thermodynamically consistent rates could be determined in practice. We analyzed the transport properties of the model out of equilibrium under varying thermodynamic driving forces, namely membrane voltages and transmembrane concentration gradients. Comparison of the antiporter model for a realistic 1:1 Na^+^:H^+^ electroneutral stoichiometry with a hypothetical 1:1 *M*^2+^:H^+^ electrogenic transport with a divalent cation as the driving ion illuminated the effect of the membrane potential on transport turnover numbers and how the presence of “leak cycles” can make it difficult to clearly distinguish electroneutral from electrogenic transport.

*Multibind* has already been used in a variety of applications. In the SAMPL6 blind prediction challenge, macroscopic p*K*_a_s for a set of small molecules were calculated from a set of independently determined standard state free energy differences between protonation states and subsequently compared to experimentally determined macroscopic p*K*_a_s (24). The inverse *multibind* approach was recently used to refine constant pH molecular dynamics data with micro-scale thermophoresis measurements to determine microscopic p*K*_a_ values of binding site residues in the Zn^2+^/H^+^ antiporter YiiP (58). The approach also shows promise in the construction of a full multiscale model of *Tt*NapA, a Na^+^/H^+^ transporter protein, from long time conventional equilibrium molecular dynamics and constant pH molecular dynamics data where we have been able to calculate turnover numbers as a function of the membrane potential and the transported and driving ion concentrations (Kenney et al., in preparation).

There is no limit to the connectivity of a kinetic network and more complex models than shown here are possible, although it becomes increasingly difficult to obtain all microscopic parameters. In this situation, the *multibind* approach makes it easier to include estimates (possibly with larger errors) for otherwise unknown parts of the system and still obtain a thermodynamically consistent model that can then be investigated. Overall, thermodynamically consistent kinetic models provide a powerful tool into probing the influence of a system’s microscopic features on its macroscopic function.

## AUTHOR CONTRIBUTIONS

OB and IMK designed research. IMK performed research, contributed analytic tools, and analyzed data. IMK and OB wrote the paper.

## ACKNOWLEDGMENTS

Research reported in this work was supported by the National Institute Of General Medical Sciences of the National Institutes of Health under Award No. R01GM118772. We are grateful to Nikolaus Awtrey for many helpful discussions.

## DECLARATION OF INTERESTS

The authors declare no competing interests.

## A APPENDIX

### A.1 Error estimate for state free energies

The errors of state free energies are estimated by the Cramér-Rao lower bound, which is computing by taking the inverse of the Fisher information matrix, **I**, for the maximum likelihood estimator, to achieve the lower bound on the covariance matrix (32, 33). The Fisher information matrix must first be calculated as which is just the expectation value for the negative Jacobian elements of the log likelihood’s gradient in parameter space. Summing over all defined connections between states *i* and *j*, each element of the Jacobian can be calculated as

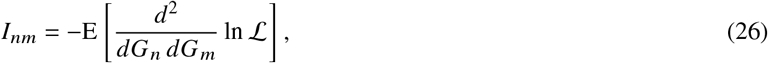

which is just the expectation value for the negative Jacobian elements of the log likelihood’s gradient in parameter space. Summing over all defined connections between states *i* and *j*, each element of the Jacobian can be calculated as

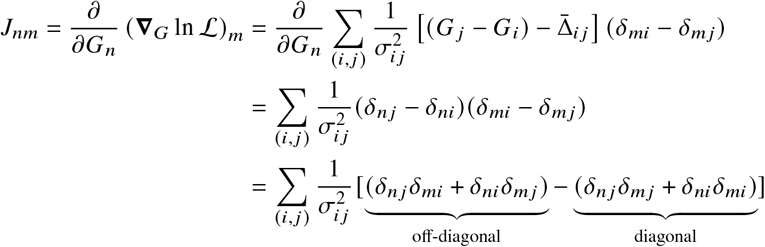

The Jacobian is a constant and hence its expectation value is itself,

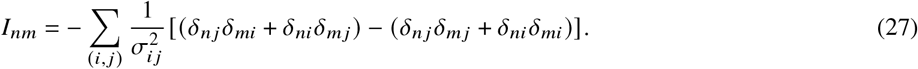

The inverse of the Fisher information, I, provides the lower bound for the covariance matrix, C, and is found with the Moore-Penrose inverse, C = I^+^. The square roots 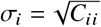 of the diagonal elements of C represent the lower bound on the standard error of each state’s free energy.

### A.2 Error estimate for effective free energy differences

Using the computed standard errors *σ*_*s*_ for the state free energies *G*_*s*_, standard errors for the effective free energy differences Δ*G*_*A,B*_ are computed as

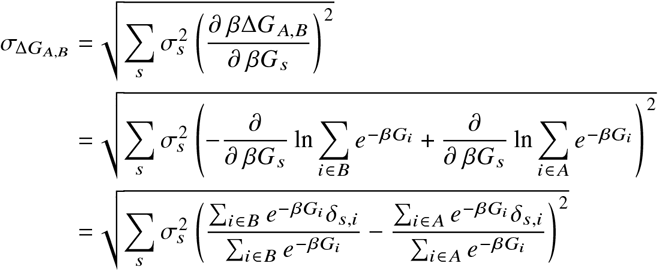

where the sum over *s* is over all states that exist in either *A* or *B*. The sums over *i* ∈ *A* or *i* ∈ *B* are over just the microstates in *A* or *B*, respectively. *δ*_*s,i*_ is the Kronecker delta, which will select the relevant terms (i.e. where state *s* is state *i*). Since one microstate cannot exist in both *A* and *B*, only one term within the parentheses remains for each term in the outer sum.

### A.3 Error of the thermodynamically consistent rates

The thermodynamically consistent rates (*k* _*ji*_ from Equation 18 and *k*_*i j*_ from Equation 19) were computed with the maximum-likelihood projection approach under the assumption of a fixed free energy difference Δ*G*_*i j*_. However, there is also an uncertainty 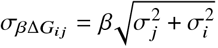 associated with the free energy difference. Both terms under the square root are from Equation 10 and for notational convenience, the free energy difference Δ*G*_*i j*_ is expressed as a reduced free energy through multiplication by *β* =(*kT*) ^−1^. Given 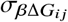, we obtain the uncertainties in the rates *k* _*ji*_ and *k*_*i j*_ by error propagation: The projection error for *k* _*ji*_ is

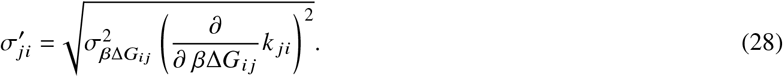

We denote this quantity with a prime to distinguish it from the uncertainty of the rate under the assumption of a fixed Δ*G*_*i j*_ (Equation 20). The error of the reverse rate, *k*_*i j*_, is

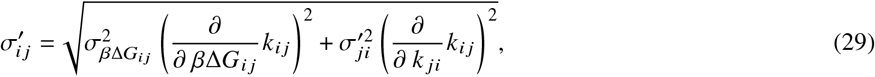

where the propagated error in the second term in the square root is the error obtained from Equation 28 (and not *σ*_*ji*_ or 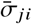). Ultimately, the errors of *k* _*ji*_ and *k*_*i j*_ depend on 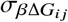, which characterizes the uncertainty of the thermodynamic consistency line (Equation 16). As this quantity goes to zero, the uncertainty of the projection (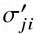 or 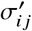) goes to zero since the line becomes well-defined, regardless of the initial errors on the forward and backward rates.

If the estimates for the free energy difference come from the maximum likelihood approach where the 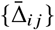 were defined by the rates, the errors of the input rates are included in the 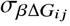 through the construction of the estimator. Then direct evaluation of the derivatives leads to standard errors for the projected rates,

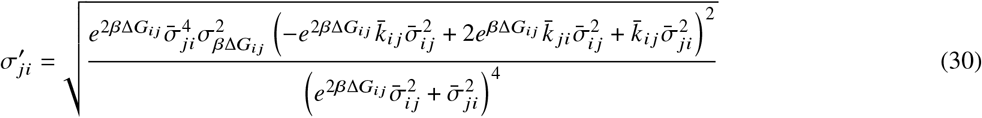

and

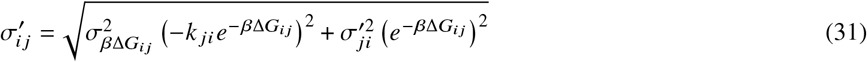

where the summary statistics of the input rates are indicated with a bar. All error estimates for thermodynamically consistent rates in this work were computed following Equations 30 and 31.

## Notes

https://github.com/Becksteinlab/multibind-publication-code

https://github.com/Becksteinlab/multibind

### Competing Interest Statement

The authors have declared no competing interest.

### Summary of Updates

(1) Added a theoretical justification for the rate projection approach as a maximum likelihood estimate with thermodynamic consistency as a constraint under the assumption of normal-distributed rates. (2) Clarified limitations of the multibind approach. (3) Discuss a leak transition as part of the antiporter model in addition to a model with ideal coupling.

