## Supplemental Information for "Thermodynamically consistent determination of free energies and rates in kinetic cycle models"

Manuscript submitted to **Biophysical** *Reports*
Article

### Supplementary Information: Thermodynamically consistent determination of free energies and rates in kinetic cycle models

Ian Kenney[1] and Oliver Beckstein[1,2]

[1]Arizona State University, Department of Physics, Tempe AZ, USA
[2]Arizona State University, Center for Biological Physics. Tempe AZ, USA

#### 1 PRACTICAL EFFECTS OF MISSING STATE DEFINITIONS

The *multibind* methods requires that all states are defined but in practice this may be difficult to achieve. The quality of the model's predictions depend on the definition of all major states so in this section we are analyzing a simple example of a hidden state to provide some insights into the limitations of the *multibind* approach when unsuitable input data are provided.

State boundaries are often difficult to define, which can affect the predicted quantities such as the free energy difference between states. Take, for instance, a three state model whose physical states are well separated into states 0, 1, and 2. Transitions are only possible between states 0 and 1, and states 1 and 2. Suppose that state 1 is difficult to measure experimentally and the effective rates (denoted by an asterisk) from state 0 to and from 2, $k^*_{02}$ and $k^*_{20}$, are calculated as the inverse mean first passage times (1). I.e., we assume that we can only measure times between being state 0 and 2 and are somehow not able to notice if the system is actually in state 1. For simplicity, we focus on equilibrium and the free energy differences between states. In particular, the input for a *multibind* model would be the measured free energy difference between states 0 and 2,

$$\beta\Delta G^*_{02} = -\ln\frac{k^*_{02}}{k^*_{20}}. \tag{S1}$$

We will now quantify under which conditions this measured free energy difference (Equation S1) accurately represents the true free energy difference, $\Delta G_{02}$. We start with the closed form expressions for the measured rates

$$k^*_{02} = \frac{k_{01}k_{12}}{k_{21}+k_{10}+k_{12}} \quad \text{and} \quad k^*_{20} = \frac{k_{21}k_{10}}{k_{01}+k_{10}+k_{12}}$$

in terms of the true underlying rates $k_{01}, k_{10}, k_{12}, k_{21}$. The effective free energy difference is

$$\beta\Delta G^*_{02} = -\ln\frac{k^*_{02}}{k^*_{20}} = -\ln\frac{k_{01}k_{12}}{k_{21}k_{10}} - \ln\frac{k_{01}+k_{10}+k_{12}}{k_{21}+k_{10}+k_{12}}$$

where the first term on the RHS is the true free energy difference, $\Delta G_{02} = -\ln\frac{k_{01}k_{12}}{k_{21}k_{10}}$, and the second term is an approximation error, $\delta_{02}$. The deviation vanishes under two conditions. In the first case, it vanishes when the sum of the rates moving out of the hidden state ($k_{10}+k_{12}$) is much larger than both of the rates going into the hidden state ($k_{01}$ and $k_{21}$), indicating that state 1 is a short-lived intermediate. In the second case, this term vanishes when $k_{01}$ and $k_{21}$ are equivalent, regardless of their relative magnitude to the other rates. From the rates we can obtain the state probabilities,

$$P_0 = \frac{k_{10}k_{21}}{k_{10}k_{21}+k_{01}(k_{12}+k_{21})} \overset{k_{01}=k_{21}}{\Longrightarrow} \frac{k_{10}}{k_{01}+k_{10}+k_{12}} \tag{S2}$$

$$P_1 = \frac{k_{01}k_{21}}{k_{10}k_{21}+k_{01}(k_{12}+k_{21})} \overset{k_{01}=k_{21}}{\Longrightarrow} \frac{k_{01}}{k_{01}+k_{10}+k_{12}} \tag{S3}$$

$$P_2 = \frac{k_{01}k_{12}}{k_{10}k_{21}+k_{01}(k_{12}+k_{21})} \overset{k_{01}=k_{21}}{\Longrightarrow} \frac{k_{12}}{k_{01}+k_{10}+k_{12}}. \tag{S4}$$

Thus, when the incoming rates to the hidden state 1 are the same, the relative populations of state 0, $P_0$, and 2, $P_2$ are determined entirely by the rates of leaving state 1 to state 0 or 2. If the incoming rates, $k_{01} = k_{21}$, are small relative to the outgoing rates, $k_{10}$ and $k_{12}$, then the hidden state is short-lived with a small population $P_1$, which is just the first case. In both cases the measured $\Delta G^*_{02}$, which serves as input for the *multibind* model, is approximately equal to the true value $\Delta G_{02}$ and thus the resulting potential graph or kinetic model will represent the underlying data well. On the other hand, if the incoming rates are relatively large, the hidden state population $P_1$ is significant, and thus the measured free energy difference $\Delta G^*_{02}$ differs from the true value and hence the resulting *multibind* model will be incorrect.

#### 2 MICROSCOPIC PROTONATION STATES OF DTPA FROM NMR DATA

##### 2.1 NMR data

NMR data for the chemical shifts of the central and terminal nitrogens of diethylenetriaminepentaacetic acid (DTPA) were extracted from Figure 4 of Submeier and Reilley (2) using the WebPlotDigitizer tool(3). The raw data are reported in Table S1 and available as CSV files from the data repository (see main paper). Each data set was extracted independently. Note that the difference in number of data points was also present in the original data.

| pH (central) | ppm (central) | pH (terminal) | ppm (terminal) |
|---|---|---|---|
| 3.1005291729704183 | 3.6928879974260154 | 3.141767697857979 | 3.9610068662417772 |
| 3.500318325266809 | 3.681291373727555 | 3.523593721119683 | 3.961438145635521 |
| 4.0082148455951305 | 3.615599991785155 | 3.9778746825304463 | 3.9468910232274763 |
| 4.588237710248704 | 3.516857547731676 | 4.304880302853974 | 3.9322001410215166 |
| 4.877619337746529 | 3.432847060112133 | 4.577503645337732 | 3.9264839776282376 |
| 5.16787721544117 | 3.3970303333173604 | 4.886381839714669 | 3.914784668359838 |
| 5.657974903646707 | 3.352403184621809 | 5.177242139419619 | 3.912101152132095 |
| 6.312259972480266 | 3.3380819704676306 | 5.68639826940586 | 3.9156883013753023 |
| 6.784942188024123 | 3.335603825379766 | 6.304592783258144 | 3.9163865632508887 |
| 7.075857253366374 | 3.335932419203571 | 6.795183362199387 | 3.898868405019271 |
| 7.494321487982364 | 3.351465323083032 | 7.086098427541637 | 3.899196998843076 |
| 7.804021166918816 | 3.384947664587855 | 7.503686411960814 | 3.866536141897767 |
| 8.022481294112008 | 3.4002546602134496 | 7.812181246876646 | 3.83375206226853 |
| 8.259123612889091 | 3.415582192953032 | 8.047892549819615 | 3.7978737241317937 |
| 8.587279311595939 | 3.4641456218295836 | 8.265640723727895 | 3.7740232890872627 |
| 8.89703375616969 | 3.5006400733859544 | 8.591277203118903 | 3.6840296555925987 |
| 9.479192480677998 | 3.5193699213428538 | 8.897964772003805 | 3.551845944262273 |
| 10.005216426952908 | 3.4506869664628934 | 9.531877023761439 | 3.41701979093218 |
| 10.512236697084413 | 3.3368018236957226 | 10.003299629647376 | 3.3452631146587075 |
| 11.601306160449624 | 3.235622308782355 | 10.493452083490212 | 3.303648076014705 |
| 12.510196677094957 | 3.224600724275554 | 11.601689519910732 | 3.256707079143192 |
| | | 12.492616907521374 | 3.2577133977285957 |

Table S1: Raw hydrogen chemical shift data of *central* and *terminal* nitrogen atoms in DTPA from 1D NMR, extracted from Figure 4 in Submeier and Reilley (2).

Each data set in in Table S1 was normalized by subtracting the minimum chemical shift from all entries and dividing by the resulting maximum.

##### 2.2 Inverse *Multibind* for NMR data

In order to constrain the *Multibind* model for DTPA we used the normalized 1D NMR shift data (based on data in Table S1) and interpreted the titration curves as site-resolved mean proton occupancy. The goodness-of-fit of the model to the experimental data was measured by the RMSD between the computed and the experimental titration curve. The RMSDs for different training data sets (only central nitrogen, only terminal nitrogens, or all central and terminal nitrogen) to the other data sets are shown in Table S2.

The microscopic p$K_a$ values of the inverse *Multibind* model were reported in the main paper when all experimental data

| Training data | $RMSD_{terminal}$ | $RMSD_{central}$ | $RMSD_{all}$ |
|---|---|---|---|
| terminal | 0.012 | **0.094** | 0.066 |
| central | **0.067** | 0.017 | 0.049 |
| all | 0.030 | 0.027 | 0.029 |

Table S2: Root-mean-square deviation (RMSD) of the calculated mean proton occupancy curves to experimental NMR data. The *Training data* indicates which data set was used for fitting the model. The RMSD to either the *terminal* or the *central* nitrogen data or the combined (*all*) data is computed. RMSD values in **bold** are considered validation as the experimental comparison data set was not used for training.

were used for training. Table S3 shows the p$K_a$ values also for the reduced training data sets (*central* and *terminal*).

| State 1 | State 2 | Central[a] | Terminal[b] | All[c] |
|---|---|---|---|---|
| 000 | 001 | 9.3 | 9.5 | 9.0 |
| 000 | 010 | 10.1 | 10.2 | 10.1 |
| 000 | 100 | 9.3 | 9.5 | 9.0 |
| 001 | 011 | 8.4 | 8.4 | 9.0 |
| 001 | 101 | 9.3 | 9.4 | 9.8 |
| 010 | 011 | 7.6 | 7.7 | 7.9 |
| 010 | 110 | 7.6 | 7.7 | 7.9 |
| 100 | 101 | 9.3 | 9.4 | 9.8 |
| 100 | 110 | 8.4 | 8.4 | 9.0 |
| 011 | 111 | 5.6 | 5.5 | 5.5 |
| 101 | 111 | 4.7 | 4.4 | 4.7 |
| 110 | 111 | 5.6 | 5.5 | 5.5 |

[a] Fit was performed only on the titration curve of the central nitrogen atom.
[b] Fit was performed only on the titration curve for the terminal nitrogen atoms.
[c] Fit was performed on the titration curves of both the central and terminal nitrogen atoms.

Table S3: Microscopic p$K_a$ values for DTPA from the inverse *Multibind* approach, split by experimental training data. States are labeled as described in the main paper.

#### 3 ANTIPORTER MODEL

The process for developing the model of a sodium/proton antiporter is shown in Fig. S1. We started by defining states, based on the inward facing (IF) and outward facing (OF) conformations and binding of individual ions (4).

##### 3.1 Model parameters

To define the equilibrium system we chose binding constants for sodium ions, measured by $K_D$, and p$K_a$s for protons (estimated parameters in Table S4). The free energy differences between the IF and OF conformations were calculated from the rates in Table 2, which had been chosen so that the free energy differences were $< 1kT$. Because we are also interested in modeling transport out of equilibrium we required forward and backward rates for ion binding processes. As these rates are often difficult to obtain, we estimated on-rates by assuming a diffusion limited process as described in section 3.3 and calculated off-rates from the equilibrium relationship

$$K_D = \frac{k_{off}}{k_{on}}. \tag{S5}$$

The rates of the model are listed in the main paper in Table 2.

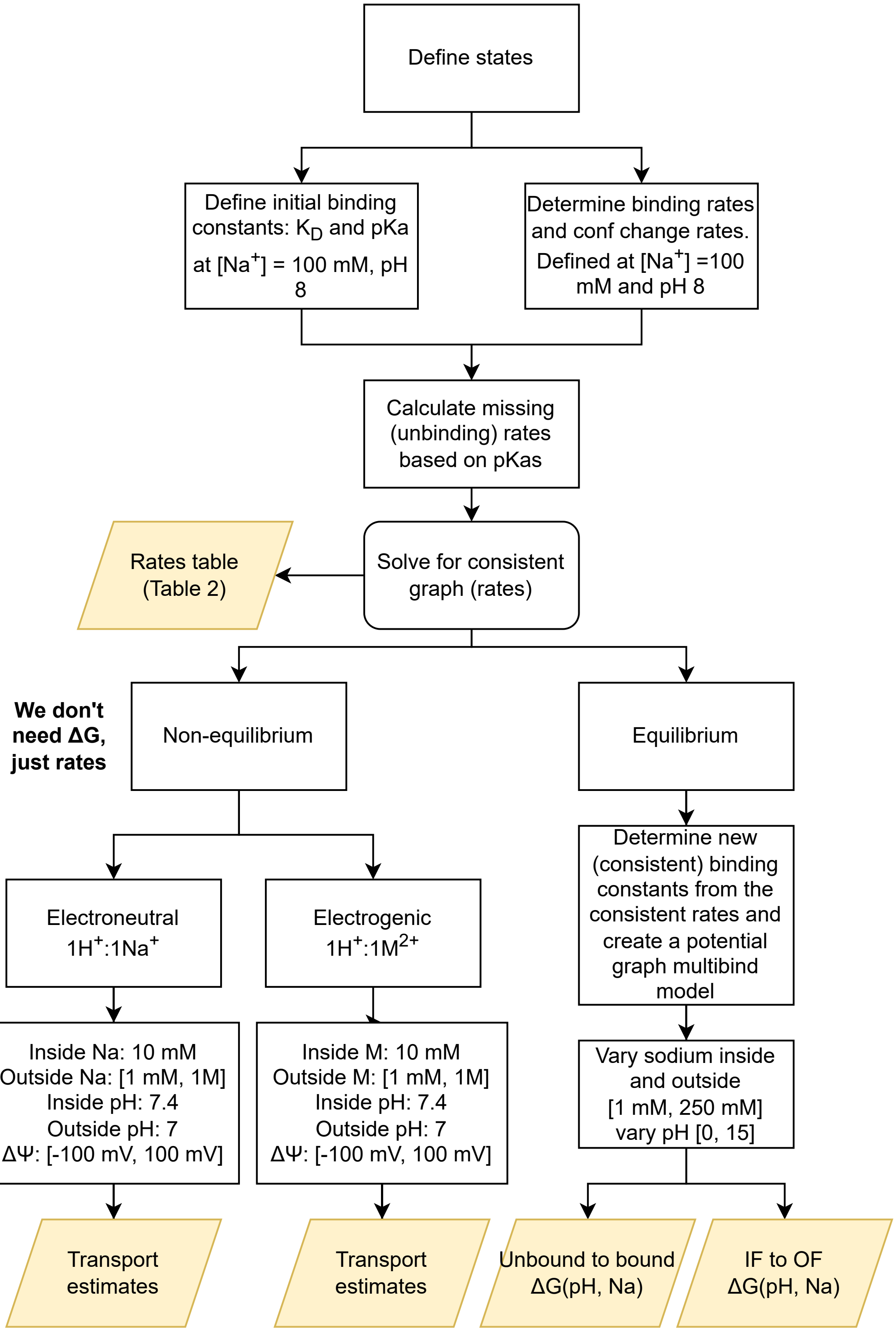


Figure S1: Flowchart following the development of both equilibrium and non-equilibrium models of a toy sodium/proton antiporter.

| conformation | $Na^+$ | $H^+$ |
|---|---|---|
| | initial $K_D$ | $pK_a$ |
| Inward | 22 mM | 6.4 |
| Outward | 53 mM | 6.5 |
| | consistent $K_D$ | $pK_a$ |
| Inward | 24.0 mM | 6.4 |
| Outward | 52.2 mM | 6.5 |

Table S4: Equilibrium binding constants ($K_D$ for sodium ions and $pK_a$ for protons) in the sodium/proton antiporter model. The initial values were calculated from the original, inconsistent, free energy differences. The consistent values were calculated once the maximum likelihood estimator determined the set of state free energies.

#### 3.2 Model analysis

Once the model was fully defined, we solved it with the *Multibind* approach as described in the main paper and obtained a set of thermodynamically consistent rates (Table 2 in the main paper and Table S5).

In order to obtain non-equilibrium steady-state transport estimates under a membrane potential, we modified the rates involving charges moving through the membrane as described in the main paper. Given the rates $k_{ij}$, the master equations (5)

$$\frac{dp_i}{dt} = \sum_{j \neq i} k_{ji} p_j - k_{ij} p_i, \qquad 1 = \sum_i p_i. \tag{S6}$$

for the changes in state populations $p_i$ were solved for the $p_i$ under steady-state conditions, $dp_i/dt = 0$, using singular value decomposition (SVD). The net transition flux from state $i$ to state $j$

$$J_{ij} = k_{ij} p_i - k_{ji} p_j \tag{S7}$$

was calculated from the steady state populations $p_i$ and $p_j$ and the transition rates $k_{ij}$ and $k_{ji}$ between states $i$ and $j$. $J_{ij} > 0$ when there is net flux from $i$ to $j$ and $J_{ij} < 0$ when the net flux is in the opposite direction; in equilibrium detailed balance $J_{ij} = 0$ holds for all net fluxes.

For the equilibrium case, the thermodynamically consistent equilibrium binding constants were calculated from the consistent rates. They were close to the initial estimated parameters (Table S4), indicating that the chosen estimates were already close to a consistent model. The equilibrium model was solved for equal inside and outside sodium concentrations and for pH ranging from 0 to 15 in order to calculate the pH-dependence of the sodium binding free energy and the free energy difference between IF and OF conformation.

#### 3.3 Diffusion limited ligand binding

For the antiporter model, order of magnitude estimates for ligand binding and unbinding rates are required. Given a free energy between the bound and unbound states, the ligand binding rate is used to calculate the unbinding rate or vise versa. With the diffusion coefficients of various ions known, we calculate the binding rate first. The binding process is treated as the diffusion of a ligand from bulk over a capturing surface. We assume that once the ligand crosses this surface binding occurs immediately. The radius of the capture surface, $R$, is set to reflect the patch size of the negatively charged surface near the binding pocket of an antiporter. The concentration of the ligand in equilibrium is treated as bulk concentration, $c_0$, far away from the capture surface and zero at the capture surface. In cylindrical coordinates, the ligand concentration must satisfy Laplace's equation

$$\frac{1}{r}\frac{\partial}{\partial r}\left(r\frac{\partial c}{\partial r}\right) + \frac{1}{r^2}\frac{\partial^2}{\partial \theta^2}c + \frac{\partial^2 c}{\partial z^2} = 0 \tag{S8}$$

| State 1 | State 2 | intrinsic rate constant and error | reaction |
|---|---|---|---|
| OF(0) | OF($H^+$) | 19446702398.8 ± 356746472.4 (M s) $^{-1}$ | OF(0) + $H^+$ $\longrightarrow$ OF($H^+$) |
| OF($H^+$) | OF(0) | 6008.0 ± 4.1 (M s) $^{-1}$ | OF($H^+$) $\longrightarrow$ OF(0) + $H^+$ |
| OF(0) | OF($Na^+$) | 3200440627.7 ± 3929.5 (M s) $^{-1}$ | OF(0) + $Na^+$ $\longrightarrow$ OF($Na^+$) |
| OF($Na^+$) | OF(0) | 167079476.7 ± 3240.2 (M s) $^{-1}$ | OF($Na^+$) $\longrightarrow$ OF(0) + $Na^+$ |
| IF(0) | IF($H^+$) | 21584542374.6 ± 407685212.2 (M s) $^{-1}$ | IF(0) + $H^+$ $\longrightarrow$ IF($H^+$) |
| IF($H^+$) | IF(0) | 7889.0 ± 30.9 (M s) $^{-1}$ | IF($H^+$) $\longrightarrow$ IF(0) + $H^+$ |
| IF(0) | IF($Na^+$) | 3199480588.0 ± 64268.0 (M s) $^{-1}$ | IF(0) + $Na^+$ $\longrightarrow$ IF($Na^+$) |
| IF($Na^+$) | IF(0) | 76760846.6 ± 2645.8 (M s) $^{-1}$ | IF($Na^+$) $\longrightarrow$ IF(0) + $Na^+$ |
| IF($Na^+$) | OF($Na^+$) | 4990.0 ± 18.5 s $^{-1}$ | IF($Na^+$) $\longrightarrow$ OF($Na^+$) |
| OF($Na^+$) | IF($Na^+$) | 8006.2 ± 12.1 s $^{-1}$ | OF($Na^+$) $\longrightarrow$ IF($Na^+$) |
| IF($H^+$) | OF($H^+$) | 8006.2 ± 12.6 s $^{-1}$ | IF($H^+$) $\longrightarrow$ OF($H^+$) |
| OF($H^+$) | IF($H^+$) | 4990.0 ± 19.1 s $^{-1}$ | OF($H^+$) $\longrightarrow$ IF($H^+$) |
| OF(0) | IF(0) | 99.7 ± 2.4 s $^{-1}$ | OF(0) $\longrightarrow$ IF(0) |
| IF(0) | OF(0) | 135.2 ± 3.8 s $^{-1}$ | IF(0) $\longrightarrow$ OF(0) |

Table S5: Thermodynamically consistent intrinsic rate constants, $k^{\circ}_{ij}$, for the antiporter model, generated by the *multibind* rate projection procedure.

subject to the following boundary conditions:

$$\begin{aligned} c &= 0; & z &= 0,\ r \le R \\ \frac{\partial c}{\partial z} &= 0; & z &= 0,\ r > R \\ c &= c_0; & z &= \infty,\ \forall r \\ c &= c_0; & r &= \infty,\ \forall z. \end{aligned}$$

The movement of the ligand follows down the concentration gradient and can be represented through the diffusion coefficient and the concentration from solving Equation S8. Integrating over the disc, the particle current through the capture surface is

$$\Phi = -\int_0^R r\,dr \int_0^{2\pi} d\theta\, D \left(\frac{\partial c}{\partial z}\right)_{z=0}, \tag{S9}$$

where

$$\left(\frac{\partial c}{\partial z}\right)_{z=0} = \frac{2c}{\pi}\frac{1}{\sqrt{R^2 - r^2}}, \tag{S10}$$

as shown by Saito (6). The resulting particle current is then

$$\Phi = 4c_0 RD, \tag{S11}$$

and the single ligand binding rate is estimated as

$$k = 4c_0 RDN_A, \tag{S12}$$

where $N_A$ is the Avogadro constant. The diffusion coefficient for free protons is taken as $9.3 \times 10^{-5}$ $cm^2/s$ (7) and $1.334 \times 10^{-5}$ $cm^2/s$ for sodium ions (8).

#### 3.4 Outer cycle driving force of a simple antiporter transport cycle

The thermodynamic driving force of the outer cycle (where one proton is exchanged for another ion), $\chi$, depends on the concentration gradients of the ions as well as the membrane potential $\Delta\Psi$,

$$\chi = \Delta\Psi\,(q_{H^+} - q_M) + kT\left(\ln\frac{[H^+]_{in}}{[H^+]_{out}} + \ln\frac{[M]_{out}}{[M]_{in}}\right). \tag{S13}$$

The first term depends on the charge difference between the transported species (proton $H^+$ and cation M) while the second term consists of the chemical potential difference, which depends on the concentrations of all species on both sides of the

| reaction | no leak | **small leak** | large leak |
|---|---|---|---|
| IF(0) $\longrightarrow$ OF(0) | 0 $s^{-1}$ | 135 $s^{-1}$ | 1350 $s^{-1}$ |
| OF(0) $\longrightarrow$ IF(0) | 0 $s^{-1}$ | 100 $s^{-1}$ | 1000 $s^{-1}$ |

Table S6: Forward and reverse values of the empty binding site conformation change rates. The "small leak" was used for the primary findings and figures in the main paper. Alternate models including the "no leak" and "large leak" conformational change rates were constructed and used to calculate turnover numbers (Figure S2, Figure S3).

| State 1 | State 2 | intrinsic rate constant and error | reaction |
|---|---|---|---|
| OF(0) | OF($H^+$) | 19445150148.1 ± 358416407.7 (M s) $^{-1}$ | OF(0) + $H^+$ $\longrightarrow$ OF($H^+$) |
| OF($H^+$) | OF(0) | 6008.0 ± 4.1 (M s) $^{-1}$ | OF($H^+$) $\longrightarrow$ OF(0) + $H^+$ |
| OF(0) | OF($Na^+$) | 3200439303.3 ± 3953.6 (M s) $^{-1}$ | OF(0) + $Na^+$ $\longrightarrow$ OF($Na^+$) |
| OF($Na^+$) | OF(0) | 167088412.3 ± 3259.9 (M s) $^{-1}$ | OF($Na^+$) $\longrightarrow$ OF(0) + $Na^+$ |
| IF(0) | IF($H^+$) | 21589117545.1 ± 412330181.5 (M s) $^{-1}$ | IF(0) + $H^+$ $\longrightarrow$ IF($H^+$) |
| IF($H^+$) | IF(0) | 7888.7 ± 31.2 (M s) $^{-1}$ | IF($H^+$) $\longrightarrow$ IF(0) + $H^+$ |
| IF(0) | IF($Na^+$) | 3199482412.5 ± 65046.1 (M s) $^{-1}$ | IF(0) + $Na^+$ $\longrightarrow$ IF($Na^+$) |
| IF($Na^+$) | IF(0) | 76739017.9 ± 2677.4 (M s) $^{-1}$ | IF($Na^+$) $\longrightarrow$ IF(0) + $Na^+$ |
| IF($Na^+$) | OF($Na^+$) | 4990.0 ± 18.5 $s^{-1}$ | IF($Na^+$) $\longrightarrow$ OF($Na^+$) |
| OF($Na^+$) | IF($Na^+$) | 8006.2 ± 12.2 $s^{-1}$ | OF($Na^+$) $\longrightarrow$ IF($Na^+$) |
| IF($H^+$) | OF($H^+$) | 8006.2 ± 12.6 $s^{-1}$ | IF($H^+$) $\longrightarrow$ OF($H^+$) |
| OF($H^+$) | IF($H^+$) | 4990.0 ± 19.1 $s^{-1}$ | OF($H^+$) $\longrightarrow$ IF($H^+$) |

Table S7: Thermodynamically consistent intrinsic rate constants, $k^{\circ}_{ij}$, for the antiporter model with no leak pathway, generated by the *multibind* rate projection procedure.

membrane. For electroneutral transport, the charge difference term is zero and the cycle driving force does not depend on the membrane potential difference. For electrogenic transport at fixed ion concentrations and pH, the driving force $\chi(\Delta\Psi)$ is a linear function of the membrane potential.

#### 3.5 Variation of leak pathway

To investigate the effects of the leak pathway in the antiporter model, we performed the turnover number calculations with a model containing no leak (Figure S2) and with a model containing a leak an order of magnitude larger than that in the main paper (Figure S3). The empty binding site conformational change rates are tabulated in Table S6.

| State 1 | State 2 | intrinsic rate constant and error | reaction |
|---|---|---|---|
| OF(0) | OF($H^+$) | 19464612022.2 ± 335486336.2 (M s) $^{-1}$ | OF(0) + $H^+$ ⟶ OF($H^+$) |
| OF($H^+$) | OF(0) | 6007.7 ± 3.9 (M s) $^{-1}$ | OF($H^+$) ⟶ OF(0) + $H^+$ |
| OF(0) | OF($Na^+$) | 3200455887.1 ± 3613.5 (M s) $^{-1}$ | OF(0) + $Na^+$ ⟶ OF($Na^+$) |
| OF($Na^+$) | OF(0) | 166976457.9 ± 2981.5 (M s) $^{-1}$ | OF($Na^+$) ⟶ OF(0) + $Na^+$ |
| IF(0) | IF($H^+$) | 21531805431.9 ± 336922380.8 (M s) $^{-1}$ | IF(0) + $H^+$ ⟶ IF($H^+$) |
| IF($H^+$) | IF(0) | 7893.0 ± 25.5 (M s) $^{-1}$ | IF($H^+$) ⟶ IF(0) + $H^+$ |
| IF(0) | IF($Na^+$) | 3199459439.0 ± 52082.8 (M s) $^{-1}$ | IF(0) + $Na^+$ ⟶ IF($Na^+$) |
| IF($Na^+$) | IF(0) | 77013041.3 ± 2147.6 (M s) $^{-1}$ | IF($Na^+$) ⟶ IF(0) + $Na^+$ |
| IF($Na^+$) | OF($Na^+$) | 4989.6 ± 18.1 s $^{-1}$ | IF($Na^+$) ⟶ OF($Na^+$) |
| OF($Na^+$) | IF($Na^+$) | 8006.4 ± 11.9 s $^{-1}$ | OF($Na^+$) ⟶ IF($Na^+$) |
| IF($H^+$) | OF($H^+$) | 8006.0 ± 12.5 s $^{-1}$ | IF($H^+$) ⟶ OF($H^+$) |
| OF($H^+$) | IF($H^+$) | 4990.4 ± 19.1 s $^{-1}$ | OF($H^+$) ⟶ IF($H^+$) |
| OF(0) | IF(0) | 999.6 ± 5.7 s $^{-1}$ | OF(0) ⟶ IF(0) |
| IF(0) | OF(0) | 1350.3 ± 8.9 s $^{-1}$ | IF(0) ⟶ OF(0) |

Table S8: Thermodynamically consistent intrinsic rate constants, $k^{\circ}_{ij}$, for the antiporter model with a large leak pathway, generated by the *multibind* rate projection procedure.

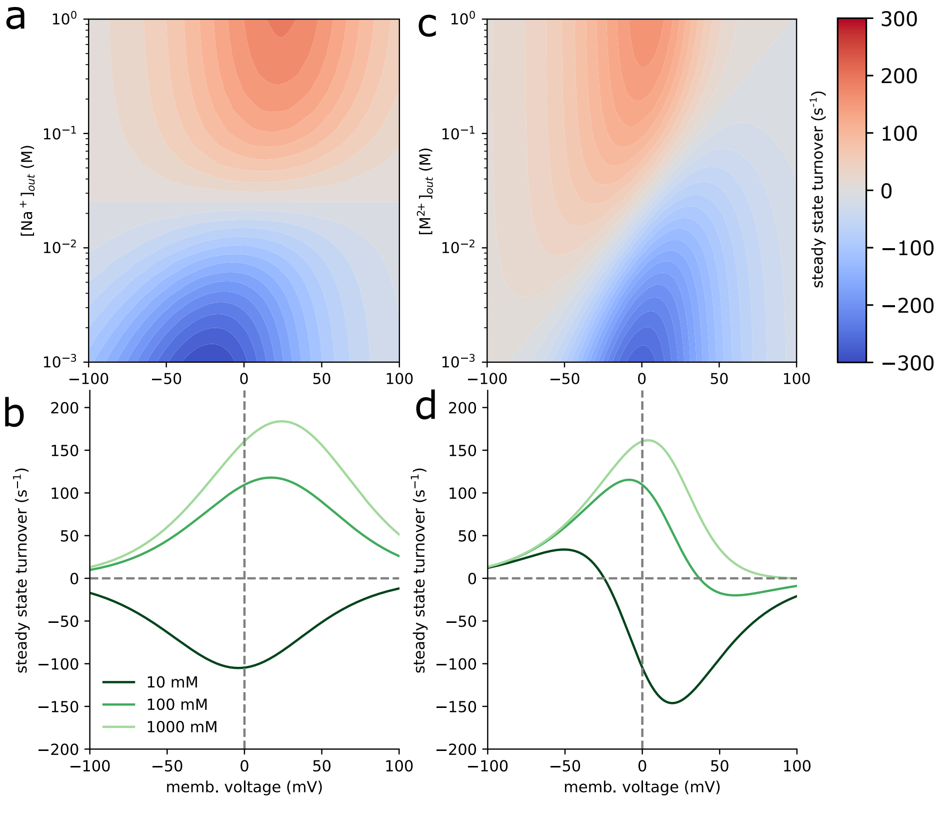


Figure S2: Non-leaking (no conformational change is possible when the binding site is empty, "no leak" column of Table S6) antiporter turnover numbers ($Na^+/H^+$ or $M^{2+}/H^+$ transport cycle completions per unit time) for the electroneutral ($Na^+/H^+$) and electrogenic ($M^{2+}/H^+$) antiporter models in steady state as a function of the cycle driving force in the driving ion gradient and an applied membrane voltage. The internal *driving* ion concentration was fixed at 10 mM for both models while the external concentration was a model parameter ranging from 1 mM to 1 M. Internal and external pH values were respectively set to 7.4 and 7. The membrane voltage, $\Delta\Psi = \Psi_{in} - \Psi_{out}$, which ranged from -100 mV to 100 mV. Turnover numbers were calculated for both the electroneutral (**a**) and electrogenic (**c**) models as a function of the external driving ion concentration and the applied membrane voltage. Turnover numbers at fixed external driving ion concentrations are shown as a function of the membrane voltage for both models (**b** and **d**).

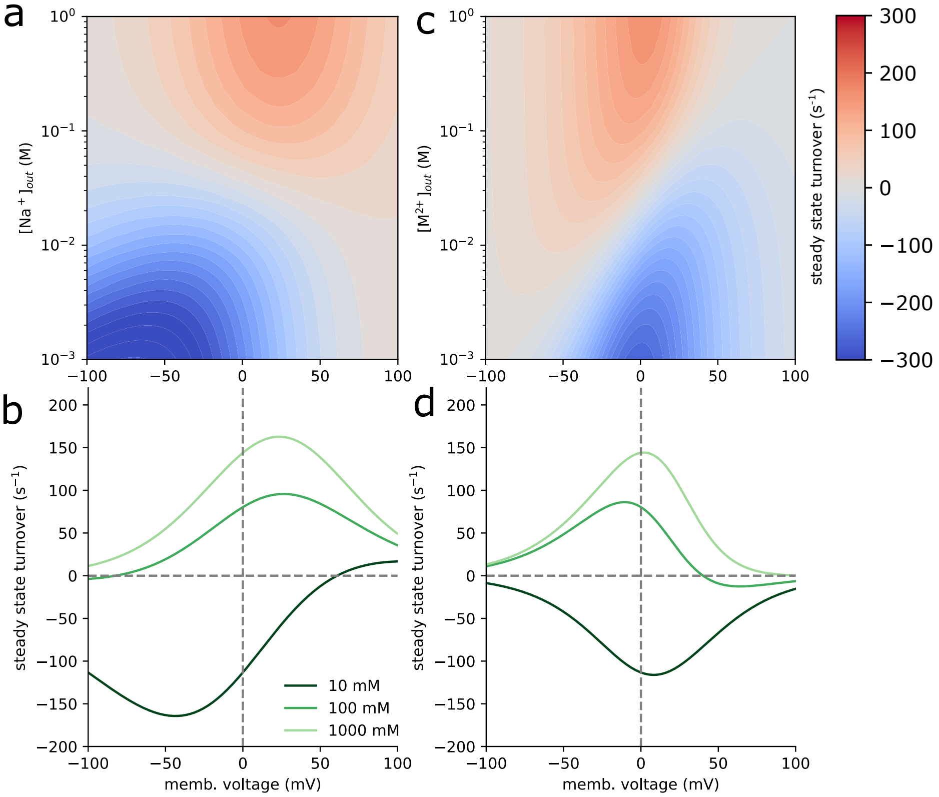


Figure S3: Leaking (transitions on the order of 1000 $s^{-1}$, "large leak" column of Table S6) antiporter turnover numbers ($Na^+/H^+$ or $M^{2+}/H^+$ transport cycle completions per unit time) for the electroneutral ($Na^+/H^+$) and electrogenic ($M^{2+}/H^+$) antiporter models in steady state as a function of the cycle driving force in the driving ion gradient and an applied membrane voltage. The internal *driving* ion concentration was fixed at 10 mM for both models while the external concentration was a model parameter ranging from 1 mM to 1 M. Internal and external pH values were respectively set to 7.4 and 7. The membrane voltage, $\Delta\Psi = \Psi_{in} - \Psi_{out}$, which ranged from -100 mV to 100 mV. Turnover numbers were calculated for both the electroneutral (**a**) and electrogenic (**c**) models as a function of the external driving ion concentration and the applied membrane voltage. Turnover numbers at fixed external driving ion concentrations are shown as a function of the membrane voltage for both models (**b** and **d**).

8. 1978. CRC Handbook of Chemistry and Physics (Online). *CRC handbook of chemistry and physics (Online)* .
